# Combination of transcriptomic, proteomic and degradomic profiling reveals common and distinct patterns of pathogen-induced cell death in maize

**DOI:** 10.1101/2022.12.23.521742

**Authors:** Sina Barghahn, Georgios Saridis, Melissa Mantz, Ute Meyer, Jaqueline C Mellüh, Johana C Misas Villamil, Pitter F Huesgen, Gunther Doehlemann

## Abstract

Regulated cell death (RCD) is crucial for plant development, as well as in decision-making in plant-microbe interactions. Previous studies revealed components of the molecular network controlling RCD, including different proteases. However, the identity, the proteolytic network as well as molecular components involved in the initiation and execution of distinct plant RCD processes, still remain largely elusive. In this study, we analyzed the transcriptome, proteome and N-terminome of *Z. mays* leaves treated with the Xanthomonas effector avrRxo1, the mycotoxin Fumonisin B1 (FB1), or the phytohormone salicylic acid (SA) to dissect plant cellular processes related to cell death and plant immunity. We found highly distinct and time-dependent biological processes being activated on transcriptional and proteome levels in response to avrRxo1, FB1 and SA. A correlation analysis of the transcriptome and proteome identified general, as well as trigger-specific markers for cell death in *Z. mays*. We found that proteases, particularly papain-like cysteine proteases, are specifically regulated during RCD. Collectivley, this study characterizes distinct RCD responses in *Z. mays* and provides a framework for the mechanistic exploration of components involved in the initiation and execution of cell death.

## Introduction

In plants, regulated cell death (RCD) plays an essential role in development, responses to environmental cues and immunity. While RCD is required for proper plant development, e.g. xylem formation and senescence, plant cell death in response to a pathogen attack can be beneficial or detrimental for the whole plant (Daneva et al., 2016; Pitsili et al., 2020). Biotrophic pathogens rely on living host tissue, while necrotrophic pathogens derive nutrients from the released content of dead cells (Pitsili et al., 2020). Thus, both, plants and microbes, have developed several strategies to induce or prevent cell death for their own benefit.

Plant immune-related RCD induced by nucleotide binding-domain leucine rich repeat containing (NLR) receptors is termed hypersensitive response (HR). HR is restricted at the site of infection and is accompanied with an elevated increase in intracellular Ca^2+^ levels as well as the generation of reactive oxygen species (ROS). The initiation of HR occurs upon the activation of NLR receptors (Pitsili et al., 2020; Saur et al., 2021; Li & Weigel, 2021). The activation of NLRs is induced by direct or indirect recognition of pathogen derived effectors and can be amplified by the recognition of microbe associated molecular patterns (MAMPs) through plasma membrane residing pattern recognition receptors (PRRs) (Zhou & Zhang, 2020; Ngou et al., 2021; Yuan et al., 2021a; Yuan et al., 2021b). NLRs can be divided into different classes according to their N-terminal domain as coiled-coiled (CC) NLRs (CNLs), TOLL/Interleukin 1 receptor (TIR)-like domain NLRs (TNLs) or RPW8-like CC domain containing NLRs (RNLs) (Maekawa et al., 2011; Jacob et al., 2013).

In the past, several NLR-effector pairs have been identified (Chen et al., 2022). The CNL Rxo1 of the monocot crop plant *Zea mays* is indispensable for the perception of the effector avrRxo1 and consequent induction of HR. AvrRxo1 is derived from the rice blast pathogen *Xanthomonas oryzae* pv. *oryzicola*, and secreted into the cell via the type III secretion system (Zhao et al., 2004; Zhao et al., 2005). Inside the cell, avrRxo1 acts as a kinase and phosphorylates NAD^+^ which is crucial for Rxo1-mediated HR induction and stops *Xanthomonas* growth (Shidore et al., 2017). Necrotrophic pathogens can secrete toxins to induce plant cell death. Examples are Fumonisin toxins, which are produced by fungal pathogens from the genus *Fusarium*. The most common one is Fumonisin B1 (FB1) which is produced by the maize pathogen *F. verticillioides* (formerly *F. moniliforme*) (Rheeder et al., 2002). FB1 disrupts the sphingolipid metabolism by effectively inhibiting ceramide synthase activity (Wang et al., 1991; Luttgeharm et al., 2016). This inhibition leads to a change in the abundance of structural and signaling sphingolipids resulting in potentially damaged structural components and the production of cell death signals. However, details of the signaling pathway induced by FB1 are not yet fully understood (Shi et al., 2007; Saucedo-García et al., 2011; Ternes et al., 2011; Kimberlin et al., 2013, Kimberlin et al., 2016; Yanagawa et al., 2017; Zeng et al., 2020).

To date, hormones, molecular components and networks that regulate and execute immune-related cell death in plants, remain largely elusive. The phytohormone salicylic acid (SA) has been shown to be involved in immunity against biotrophic pathogens and was proposed to accumulate around pathogen infection sites in a concentration gradient where it triggers various defense responses (Dorey et al., 1997; Enyedi et al., 1992; Fu et al., 2012; Yan & Dong, 2014). SA is also involved in the regulation of plant RCD, including HR, however it is not clear how SA mechanistically controls RCD (Radojičić et al., 2018). Responses towards the phytohormone jasmonic acid (JA) are induced in cells surrounding SA-active cells (Betsuyaku et al., 2018) indicating that both, SA and JA do play a role in immune-related cell death in plants. Only a few cell death-indicating transcriptional markers have been identified in dicot plants, so far. A meta-analysis of transcriptome data revealed transcriptional markers for developmental cell death in *Arabidopsis thaliana* (Olvera-Carrillo et al., 2015). In tobacco, the gene hypersensitivity-related (hsr) 203J was identified as useful cell death marker. However, hsr203j was not only induced in response to pathogens but also upon treatment with other cell death inducing agents (Pontier et al., 1994, 1998). Recently, a transcriptome analysis in *A. thaliana* comparing gene expression in cells undergoing cell death and cells of surrounding tissue revealed 13 reliable transcriptional HR markers including an uncharacterized AAA-ATPase (Salguero-Linares et al., 2022). The molecular chaperone complex consisting of SGT1, RAR1 and HSP90 was shown to interact with several NLRs and is essential for NLR-dependent immune responses in various plant-pathogen interactions (Shirasu, 2009; Zhang et al., 2010). HR is not induced if either SGT1, RAR1 or HSP90 are mutated or silenced (Liu et al., 2004; Zhang et al., 2004; Scofield et al., 2005; Azevedo et al., 2006; Shibata et al., 2011; Kim et al., 2018). Also, calmodulin or calmodulin-like proteins are required for a proper HR in *A. thaliana* and Rp1-D21-mediated HR in *Zea mays* (Chiasson et al., 2005; Ma et al., 2008; Murphree et al., 2020).

Although different proteins have been shown to be implicated in the regulation of cell death in plants, it is well known from the animal field that proteases can play an important role in the initiation and execution of RCD (Maekawa et al., 2022). In mammals, cysteine proteases of the caspase family regulate cell death pathways such as apoptosis and necroptosis (Bateman et al., 2021). However, various plant proteases of different families, including both cysteine and serine proteases, have been implicated in immune-related cell death in different pathosystems (Sueldo & van der Hoorn, 2017; Salguero-Linares & Coll, 2019). For example, the papain-like cysteine protease (PLCP) Cathepsin B positively regulates HR in tobacco and *A. thaliana* (Gilroy et al., 2007; McLellan et al., 2009; Ge et al., 2016). Furthermore, *A. thaliana* metacaspases were shown to act either as positive or negative regulators of plant HR in response to bacteria (Coll et al., 2010; Watanabe & Lam, 2011). In maize, the activity of PLCPs is linked with SA-signaling via the release of the signaling peptide Zip1 (van der Linde et al., 2012; Ziemann et al., 2018) was was also found to be implicated in 10-OPEA-induced cell death (Christensen et al., 2015).

In this study, we analyzed temporal changes in the transcriptome, proteome and N-terminome of *Z. mays* leaves in response to two distinct cell death triggers, the bacterial effector avrRxo1 and the mycotoxin FB1 in *Z. mays* leaves. RNA-sequencing revealed specific transcriptional reprogramming during immune-related cell death. Responses between the two cell death triggers were remarkably different, with only modest overlapping regulation of genes primarily related to biotic defence and immune responses. Proteome analysis showed generally smaller changes, of which many, but not all, reflected transcriptional responses. Combination of these datasets identified cell death markers that are either induced by both or one of the cell death triggers. N-terminome analysis further revealed altered protease activities, including the activation an RD19A-like cysteine protease, in response to cell death. Taken together, our data constitute a new large-scale resource of molecular changes during immune-related cell death in maize and reveal new protein markers and components potentially involved in plant regulated cell death.

## Results

### Experimental setup to investigate distinct cell death responses in maize leaves

To study the cellular processes that are activated upon cell death induction, two triggers of regulated cell death in maize leaves were used: firstly, an Avr-R dependent cell death was set-up, using the bacterial pathogen *Xoo* PXO99A carrying the Avr effector kinase avrRxo1 which is recognized by the R protein Rxo1 (Zhao et al., 2004, Zhao et al., 2005). Two controls were used: *Xoo* PXO99A transformed with an empty vector (pHMI) and *Xoo* PXO99A carrying the kinase-inactive avrRxo1 D193T (pHMI-avrRxo1 D193T), which is known to avoid Rxo1-mediated cell death (Shidore et al., 2017, Fig. 1A). Secondly, cell death was induced by the mycotoxin FB1. The phytohormone SA was included as a comparison. Upon vacuum-infiltration of whole leaves, fluorescent images of infiltrated leaves were taken over time and cell death was quantified based on autofluorescence. Besides, macroscopic pictures were taken over time and compared to fluorescent images. Microscopic cell death lesions started to appear at 48 hpi in response to FB1 and avrRxo1 and progressed over time, which is reflected in an increase in fluorescence (Fig. 1B, C, D; Fig. S1). To obtain a temporal insight into cell death processes, samples were collected at 12, 24, and 48 hpi and subjected to transcriptome analysis via RNASeq. For proteome and N-terminome analysis, samples were taken at 24, 48, 60 and 72 hpi (Fig. 1E).

**Figure 1.**
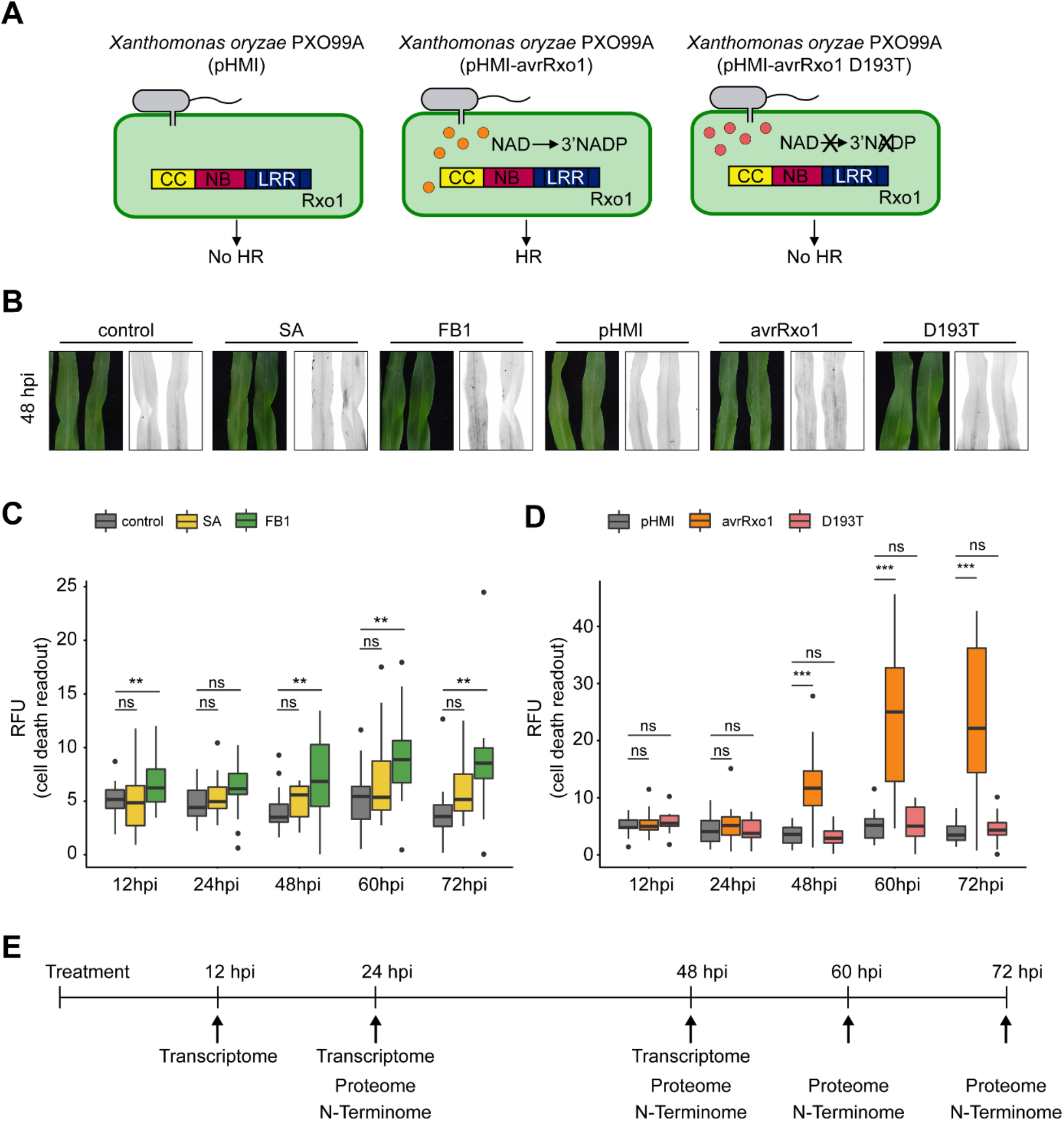
The mycotoxin FB1 and the bacterial effector avrRxo1 induce cell death in maize leaves. (A) Bacterial system to study HR in maize. Maize leaves were vacuum infiltrated with the non-host bacterial strain Xanthomonas oryzae pv. oryzae PXO99A carrying either an empty vector (pHMI, left), a vector encoding for the kinase-active effector avrRxo1 (pHMI-avrRxo1, middle) or the kinase-inactive effector avrRxo1 D193T (pHMI-avrRxo1 D193T, right). HR is induced in the presence of avrRxo1 but not the empty vector or avrRxo1-D193T. (B) Analysis of cell death development in maize. Maize leaves were vacuum-infiltrated with a control solution (DMSO), SA [2 mM], FB1 [50 μM] or Xanthomonas oryzae pv. oryzae PXO99A carrying either the empty vector (pHMI), avrRxo1 or avrRxo1 D193T (D193T) with OD600= 0.02. Colorimetric pictures were taken and a green light filter was used to detect cell death signals at 48 hpi. Cell death is indicated by black dots and is marked with an arrow. (C) and (D) Quantification of green light signals. The intensity of the green light signal (relative fluorescence units [RFUs]) was quantified using ImageLab. Boxplots show the quantification of RFUs of three to four biological replicates with each 4 leaves. Asterisks indicate significance of the treatments to the respective controls with not significant (ns) = p > 0.5, * = p ≤ 0.1, ** = p ≤ 0.05 and *** = p ≤ 0.001 calculated with an unpaired two-tailed Student’s t-test. Black dots indicate outliers. (E) Time scale and experimental design of the study. Maize leaves were vacuum-infiltrated with the same treatments as shown in B. Samples were collected at five different time points: 12, 24, 48, 60 and 72 hpi. Three biological replicates with each four leaves for 12, 24 and 48 hpi were subjected to transcriptome analysis (RNASeq). Four biological replicates with each four leaves collected at 24, 48, 60 and 72 hpi were subjected to quantitative proteome and N-terminome analysis.

### Transcriptional responses of maize leaves during cell death

We first assessed the transcriptional responses in maize leaves over time after induction of cell death with avrRxo1 and FB1. At 12 hpi, principal component analysis (PCA) clearly separated the 12 hpi samples from the other time points, but could not separate treatments. The later timepoints, at 24 hpi and 48 hpi, clustered together but showed a clear separation of the cell death triggered by avrRxo1 and FB1 samples from the rest of the samples (D193T, pHMI, SA and control) at 48 hpi (Fig. 2A). A differential gene expression analysis was performed using edgeR (Robinson et al., 2009). Genes with absolute log_2_FC>1 and FDR<0.05 were considered as significant differentially expressed genes (DEGs). At 12 hpi, the total number of DEGs was relatively similar in the avrRxo1, FB1 and D193T samples (662, 507 and 729, respectively), while SA revealed a remarkably stronger transcriptional response at this time-point, reaching more than triple DEGs (1812) (Fig. 2B). Both, avrRxo1 and FB1 showed an increasing number of DEGs at 24 and 48 hpi, with avrRxo1 tripling and FB1 doubling their DEGs at 48 hpi (2142). On the contrary, D193T and SA showed similar patterns with a decreasing number of DEGs over time (Fig. 2B).

**Figure 2:**
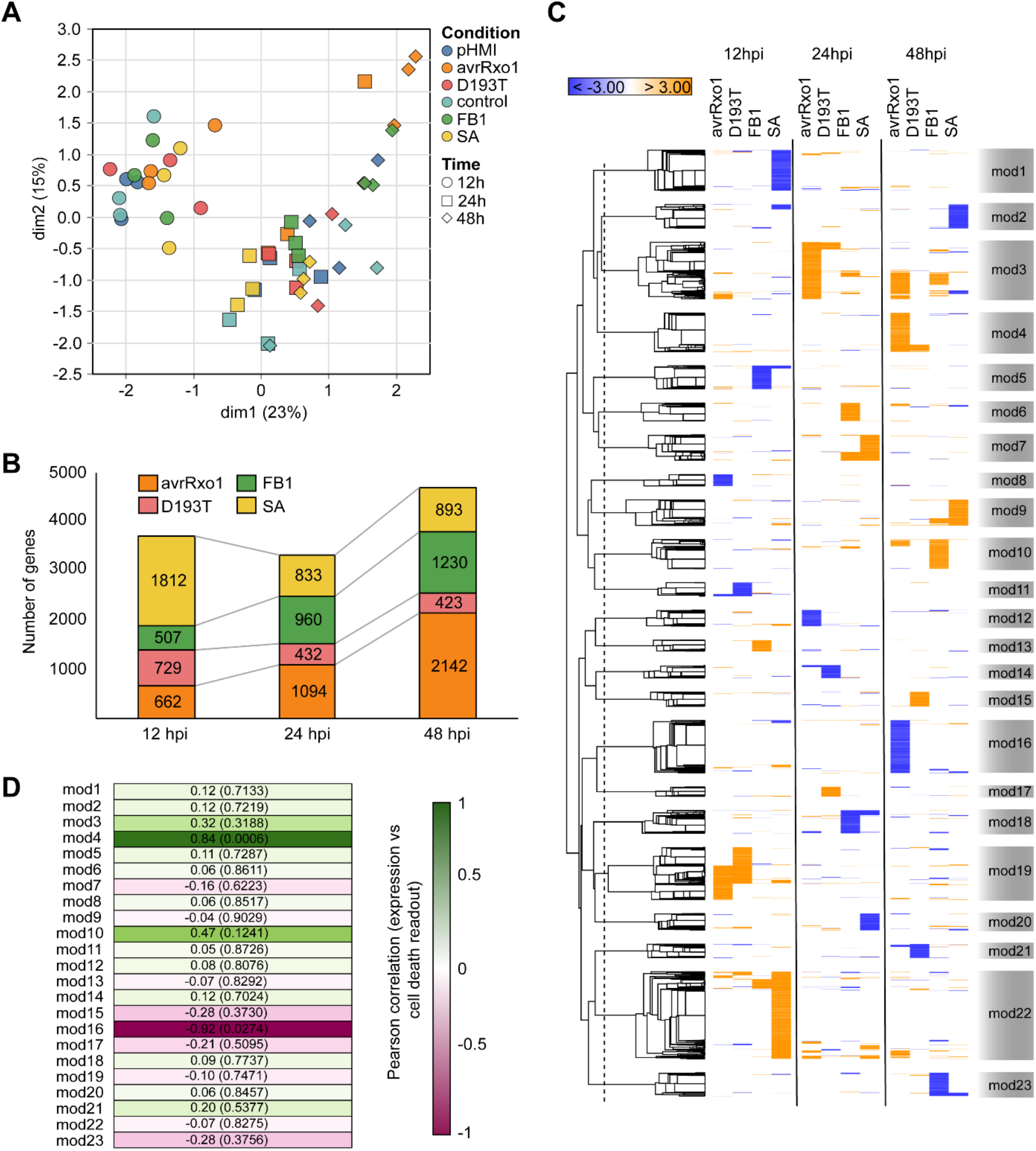
Transcriptional changes upon cell death in maize leaves. (A) MDS plot depicting the RNAseq sample distribution. Row counts were used for the calculation of the contributions of the PCs. (B) Numbers of DEGs (absolute log2FC>1, FDR<0.05). (C) Modules of co-expressed genes (only higher stringency genes, with log2FC>2 and FDR<0.05, are plotted). The metric for the distances for the hierarchical clustering was 1-r, where r is the Pearson correlation coefficient. The modules are named from 1 to 23. Scale bar represents the expression pattern (D) Pearson correlation of the gene expression pattern of each module (calculated as the average of the module’s log2FC for each condition) to the cell death readout shown in Fig. 1 C-D. Numbers show the Pearson correlation with the cell death readout and the P-values in parentheses. Correlation was considered significant if absolute correlation coefficient>0.5 and P-value<0.05.

To understand the relation of the transcriptional changes to the cell death phenotype, DEGs were hierarchically clustered (using a higher stringency setup, with absolute log_2_FC>2, to increase the sensitivity of the method) based on their expression profiles across the treatments and time-points (Fig. 2C). This clustering revealed 23 modules with distinct expression patterns (Fig. 2C, Table S1). Among these, module 4 (271 genes) and module 16 (367 genes), summarized opposite patterns of genes up- or down-regulated at 48 hpi upon avrRxo1 treatment; module 13 (78 genes), module 6 (126 genes) and module 10 (204 genes), with transcripts mainly up- regulated upon FB1 treatment at 12, 24 and 48 hpi, respectively while module 5 (163 genes), module 18 (161 genes) and module 23 (163 genes) showed mostly down-regulated genes upon FB1 treatment. Module 1 (282 gens) and module 22 (607 genes) represented genes down- or up-regulated, respectively, upon SA treatment at 12 hpi, and module 3 (396) which consisted of genes up-regulated primarily upon avrRxo1 at 24 hpi and upon both, avrRxo1 and FB1, at 48 hpi (Fig. 2C). The avrRxo1-specific module 4, which includes the aspartyl-protease Zm00001eb154620 (Table S1), showed a significant positive correlation (Pearson coefficient = 0.84, *P*-value = 0.0006) with the cell death phenotype, (Fig. 2D). In addition, a significant negative correlation (Pearson coefficient = −0.92, *P*-value = 0.0274) was observed for module 16, which contains, among others, the subtilisin-like protease Zm00001eb285310 (Fig. 2D, Table S1). These two modules indicate genes up- and down-regulated exclusively during the course of a HR-triggered response caused by the avrRxo1-Rxo1 interaction.

Most DEGs were unique to their treatments, with only very small total overlaps at each tested time point (Fig. 3A, Table S1). Some significant pair-wise overlaps were observed, namely between avrRxo1 and FB1 at 24 hpi (15% of the avrRxo1 and 19% of the FB1 up-regulated genes) and particularly at 48 hpi (31% of the avrRxo1 and 39% of the FB1 up-regulated genes). To dissect the trends of the uniquely responsive genes, the proportion of the unique up- or down-regulated genes was calculated relative to the total number of unique genes. At 12 hpi, unique DEGs were mainly represented by SA responsive genes (56% for the up-regulated set and 59% for the down-regulated set), which was dramatically reduced to below 20% (up-regulated) or 30% (down-regulated) at 24 and 48 hpi (Fig. 3B, Table S1). In contrast, avrRxo1 and FB1 treatments showed the opposite behavior, with the tendency of their unique DEGs increasing both, at the up- and down-regulated sets (Fig. 3B, Table S1). These results correlate with the general trend observed for the total number of DEGs (Fig. 2B).

**Figure 3:**
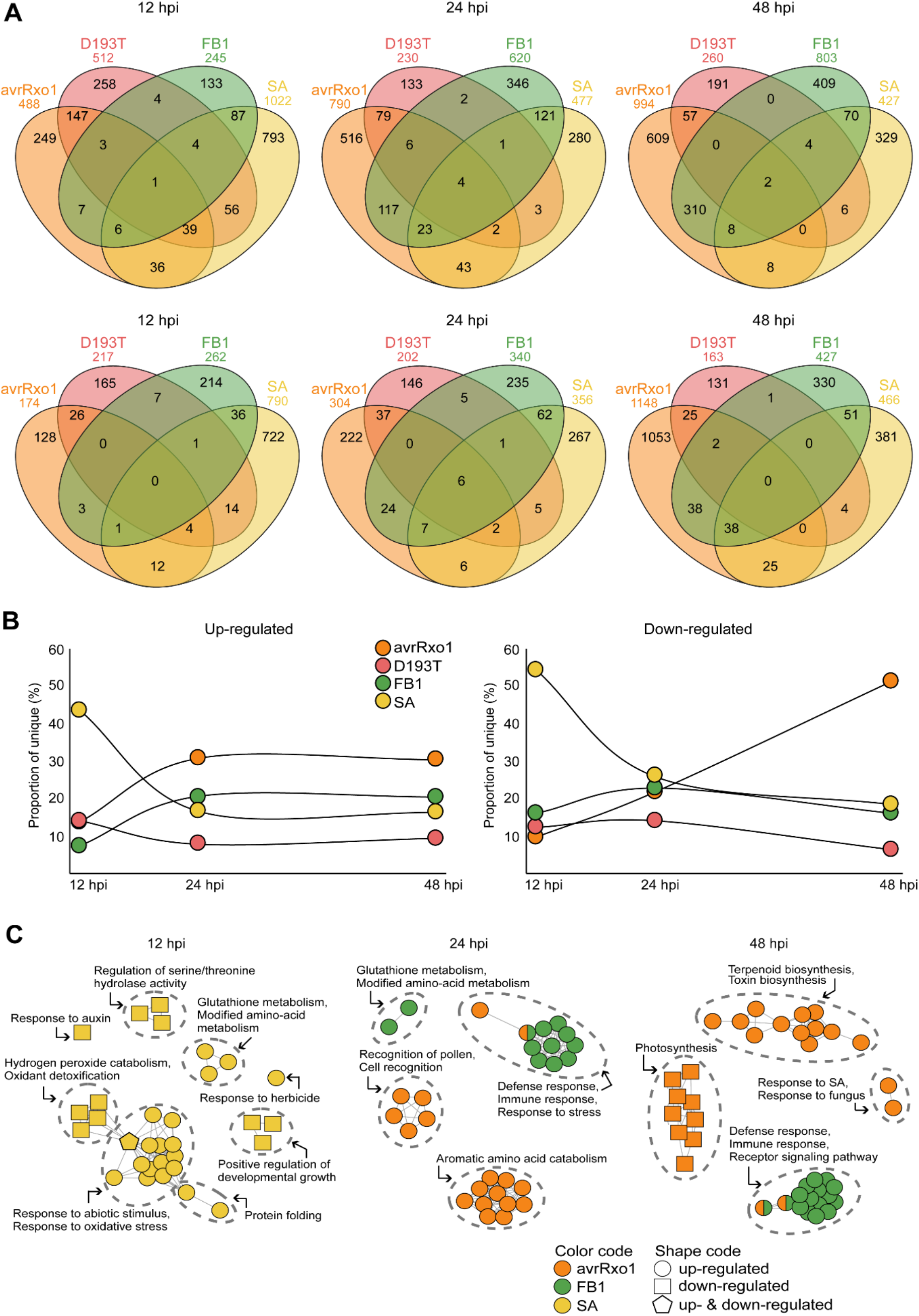
Functional characterization of the transcriptional responses upon maize leaf cell death. (A) Venn diagrams visualizing the overlap of the significantly up- (upper panel) or down-regulated (lower panel) DEGs in each treatment and time point (DEGs with absolute log2FC>1 and FDR<0.05). (B) Percentage of unique DEGs from each treatment during the time course of the experiment (C) Network analysis of enriched GO terms. GO terms enrichment was performed using a custom background including only genes expressed in the tested conditions. Only significantly enriched GO terms (FDR<0.05) were plotted and then manually curated, highlighting cluster representatives.

All DEGs from each treatment and time point were subjected to a GO enrichment analysis, with the aim of unraveling the molecular pathways represented by the detected transcriptional responses. At 12 hpi, only the SA-related gene groups showed significant enrichment of specific GO terms, as also expected from the total number of DEGs (Fig. 3C). The up-regulated pathways were associated with glutathione and modified amino-acid metabolisms, responses to abiotic stimuli and oxidative stress, as well as protein folding. In contrast, responses to auxin, the regulation of serine-threonine hydrolases and hydrogen peroxide catabolism, were among the over-represented pathways in the down-regulated gene set. At 24 hpi, FB1 treated leaves showed an over-representation of glutathione and modified amino-acid metabolisms as well as defense and immune responses. AvrRxo1 up-regulated gene sets included also defense and immune responses but also other unique GO terms were associated with cell-recognition processes and aromatic amino-acid catabolism (Fig. 3C, Table S2). At 48 hpi, both avrRxo1 and FB1 enriched GO terms included defense and immune responses, as well as receptor signaling pathways. Response to SA and terpenoid/toxin biosynthesis were among the unique GO terms enriched in the up-regulated gene set of avrRxo1 at 48 hpi, while photosynthesis-related processes were enriched in the down-regulated gene sets.

This data suggests that at the transcriptional level, SA causes rapid and unique responses typically observed at 12 hpi, up-regulating plant protein metabolisms and oxidative stress responses, while repressing auxin-related responses. On the contrary, the cell death inducing factors trigger a transcriptional reprogramming of maize leaves later on at 24 and 48 hpi, in a gradually increasing manner, maintaining a core defense- and immune-related response, but also distinctive cell death responses with protein catabolism and terpenoid biosynthesis being enriched in avrRxo1.

### Distinct proteome responses of maize leaves to cell death triggers

Next, we asked how transcriptional changes are translated to the protein level during regulated cell death in maize. To this end, we compared by quantitative proteome analysis the two cell death triggers with their respective controls, namely avrRxo1 to D193T and pHMI, and FB1 compared to SA and control treatment at four time points (24, 48, 60 and 72 hpi) (Fig. 1B). The two experiments avrRxo1/D193T/pHMI and FB1/SA/control were processed and analyzed separately using a label-free data-independent acquisition strategy, which identified 5116 proteins in the avrRxo1-experiment and 4437 proteins for the FB1 experiment, respectively. Similar to the transcript data, principal component analysis could not separate the different samples based on changes on the proteome level at the first analysed time point 24 hpi (Fig. 4A). Subtle separation appeared at 48 and was more prominent at 60 and 72 hpi for both, avrRxo1 and FB1, and separated from the rest across the dim1 (explaining 16% and 14% of the variation for avrRxo1 and FB1, respectively). However, abundance changes on the proteome level were small compared to transcriptional changes. To amplify these differences and determine differentially abundant proteins (DAPs), focusing on the protein abundance pattern across the tested time points, protein intensities were subjected to per-peptide scaling normalization. Proteins with *P*-value < 0.05, with no log_2_FC as cut-off, were considered as significantly altered in abundance in a given treatment and at a given time point. Similar to the transcriptome, at the earliest time point, at 24 hpi, the number of DAPs was similar in the avrRxo1, FB1 and D193T samples (82, 86 and 81 respectively), while SA revealed a remarkably stronger response at this time-point with almost twice as many DAPs (153) as the other treatments (Fig. 4B). However, avrRxo1-triggered cell death showed an increased number of DAPs at 48 hpi (218), which remained stable at 60 hpi (221), until it slightly dropped at 72 hpi (154). Remarkably, FB1 showed a burst of DAPs at 60 hpi (390), followed by a rapid decline at 72 hpi (156) (Fig. 4B).

**Figure 4:**
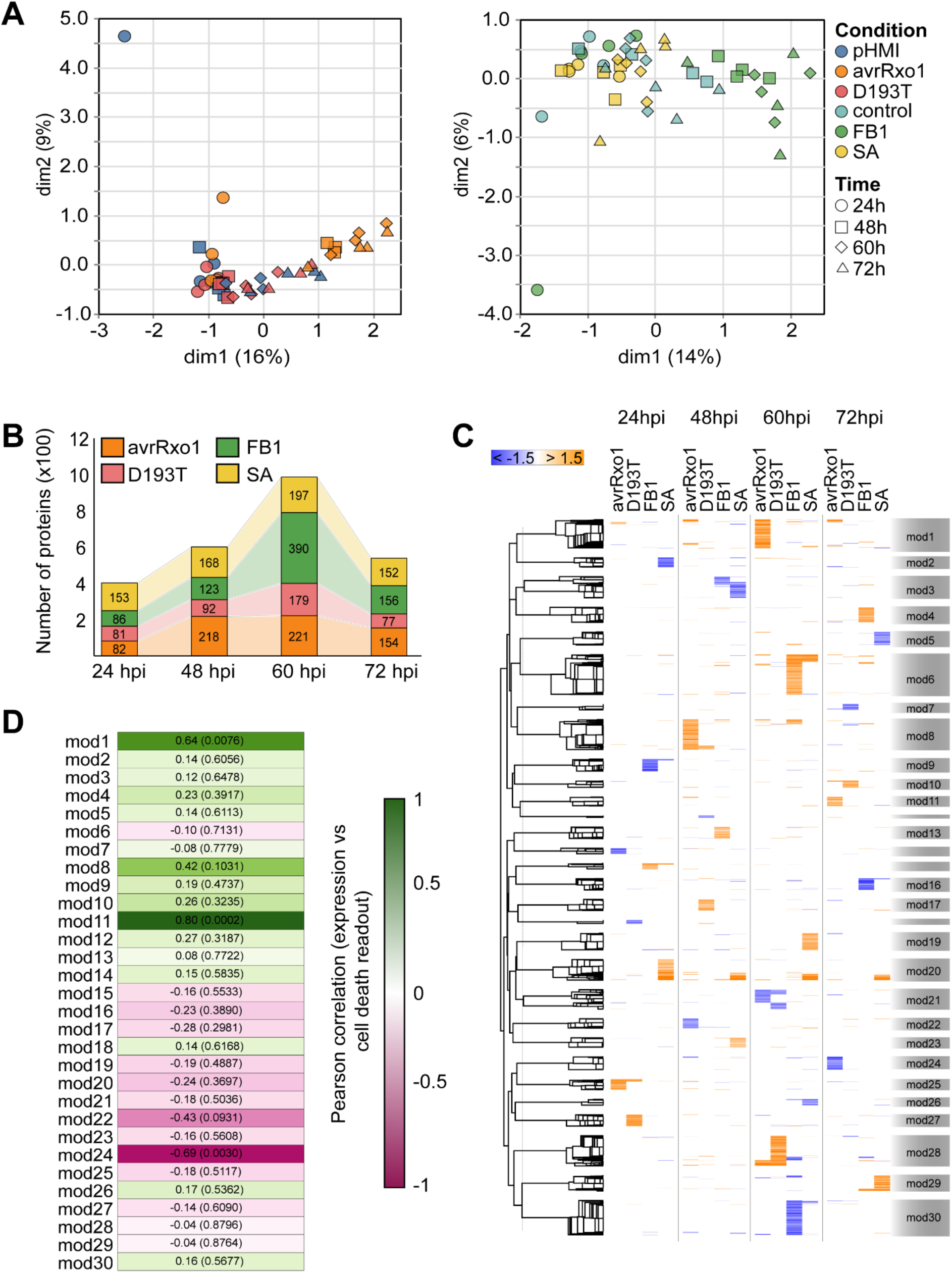
Proteome changes during cell death in maize leaves. (A) PCA plots of the avrRxo1/D193T/pHMI (left) and FB1/SA/control (right) depicting the distribution of proteomic samples. Row values were used for the calculation of the contributions of the PCs. (B) Numbers of differentially abundant proteins (P-value<0.05) over time. (C) Modules of co-accumulated proteins (only significant values are plotted, P-value<0.05). The metric for the distances for the hierarchical clustering was 1-r, where r is the Pearson correlation coefficient. Modules are named from 1 to 30. The values used for the heatmap correspond to the log2 ratios after per-row scaling normalization. (D) Pearson correlation of the pattern of the protein abundance of each module (calculated as the average of the module’s log2 ratio for each condition) to the cell death readout in Fig. 1C-D. Numbers show the Pearson correlation with the cell death readout and the P-values in parentheses. Correlation was considered significant if absolute correlation coefficient>0.5 and P-value<0.05.

Similar to the transcriptome, DAPs were hierarchically clustered based on their protein abundance profiles across the treatments and time-points, resulting in 30 distinct modules (Fig. 4C, Table S3, S4). Proteins accumulated upon avrRxo1 treatment at 24, 48, 60 and 72 hpi were found in modules 25 (41), 8 (116), 1 (110) and 11 (35), respectively, while modules 14 (22), 22 (37) and 24 (50), showed the opposite pattern with a decrease in protein abundance. Modules 15 (22), 13 (45), 6 (154) and 4 (58) typically represented FB1-accumulated proteins across the time points, while modules 9 (47), 30 (133) and 16 (44) showed FB1-decreased protein abundance. The modules 2 (37) and 20 (80) comprised of proteins depleted or accumulated upon SA treatment at 24 hpi, respectively (Fig. 4C). Compared to the transcriptome, only the avrRxo1-specific modules revealed a significant correlation to the cell death readout. More specifically, modules 1 (cor. coefficient = 0.64, P-value = 0.0076) and 11 (cor. coefficient = 0.80, P-value = 0.0002) depicted a positive correlation, while module 24 (cor. coefficient = −0.69, P-value = 0.0030), showed a negative correlation (Fig. 4D). Module 1 included three proteases (a cysteine protease, Zm00001eb066830_P001, a nudix hydrolase, Zm00001eb272540_P001 and a mitochondrial CLP protease regulatory subunit Zm00001eb182540_P001) abundant in avrRxo1 samples at 60 hpi, while 4 other avrRxo1-specific proteases (two alpha/beta hydrolases, Zm00001eb099580_P001 and Zm00001eb160350_P001, one aspartyl-protease, Zm00001eb269230_P001 and one ubiquitin-like-specific protease, Zm00001eb367170_P001), were found in module 11, enriched at 72 hpi (Table S3). Two depleted proteases were also identified in module 24, namely, the subtilisin-like protease SBT5.3, Zm00001eb235950_P001 and a glycosyl-hydrolase, Zm00001eb273940_P001 (Table S3). The overlap among DAPs in different treatments at each time point was very small. Only 12 proteins overlapped between avrRxo1 and FB1 (5.5% of avrRxo1 9.7% of FB1 DAPs) at 48 hpi, which increased to 18 (8.1% of avrRxo1 and 4.6% of FB1 DAPs) at 60 hpi, and dropped again to 8 (5.2% of avrRxo1 and 5.1% of FB1 DAP) at 72 hpi (Fig. 5A and 5B, Table S3). To further understand the pathways enriched in our lists of DAPs, all DAPs were subjected to GO enrichment analysis. Shared responses across the time points included the following: At 24 hpi, accumulated proteins represented protein biosynthesis and glutathione metabolism, in both SA and avrRxo1 (Fig. 5A). At 48 hpi, a common response between avrRxo1 and FB1 included an accumulation of proteins related to defense responses and secondary metabolism, while carboxylic acid and lipid metabolisms were found to be enriched in SA as well (Fig. 5A). During the same time, sugar and alcohol metabolism-related proteins were less abundant in both, SA and avrRxo1 (Fig. 5A) whereas protein abundance related to photosynthesis, chloroplast reorganization and autophagy decreased in response to SA and FB1. At 60 hpi, all treatments showed an over-representation of DAPS related to protein metabolism, while carbohydrate metabolism was enriched in depleted DAPs. Moreover, mitochondrial processes and defense responses were enriched among accumulating DAPs for avrRxo1 and FB1. At 72 hpi, levels of proteins involved in carboxylic acid and protein metabolisms decreased in both DAPs of avrRxo1 and FB1 (Fig. 5B). We could also observe uniquely enriched pathways for each trigger. The avrRxo1 treatment showed higher protein accumulation related to mRNA processing and ROS response at 24 hpi, accumulated protein folding and hydrolase activity at 48 hpi, and biosynthesis of small molecules at 72 hpi. In contrast, response to Cd-ions was depleted at 48 hpi, as well as proteins related to alcohol catabolism at 60 hpi. The FB1 treatment caused an accumulation of proteins related to glycerol and ether metabolisms at 48 hpi, enhanced protein transport at 60 hpi and enriched proteolysis and protein transport at 72 hpi. In contrast, a decrease in proteins of nucleotide metabolism and peptidyl-proline protein modification, as well as chloroplastic RNA and protein metabolisms at 60 hpi was observed. Unique SA-accumulating DAPs represented mitochondrial processes and gene expression, as well as S-compound and glutathione metabolisms across all time points, while unique SA-depleted DAPs corresponded to detoxification-related pathways, especially at later time points.

**Figure 5:**
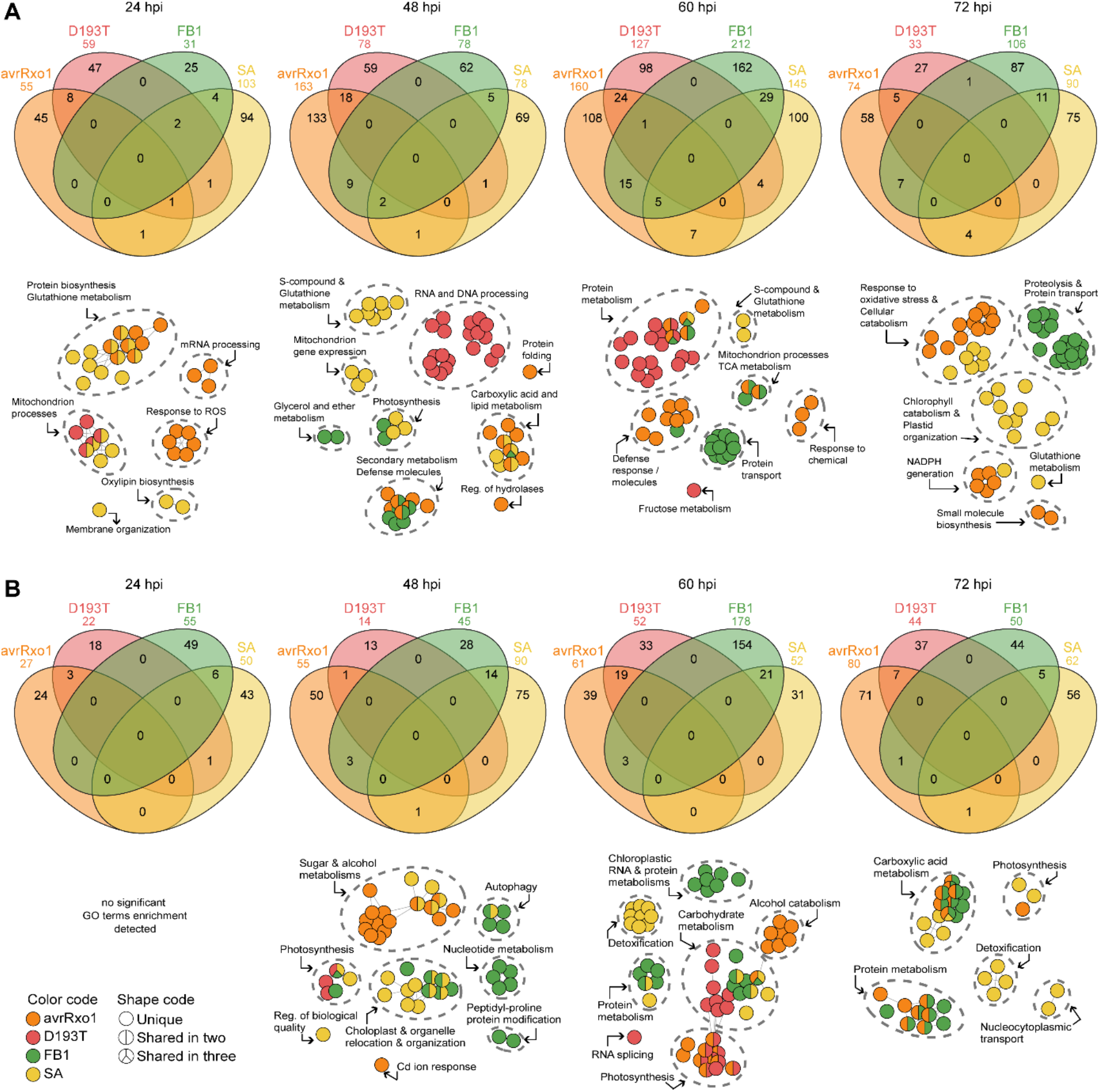
Functional characterization of proteomic responses upon maize leaf cell death. Venn diagrams visualizing the overlap of the significantly accumulating (A) or depleted (B) proteins in each treatment and time point (P-value<0.05). Accompanying networks represent enriched GO terms in significantly accumulating (A) and depleted (B) protein sets per time point. Different colors represent the different treatments. GO term enrichment was performed using a custom background including only proteins detected in our dataset. Only significantly enriched GO terms (FDR<0.05) were plotted and then manually curated highlighting cluster representatives.

In summary, the proteome data showed the strongest changes in response to cell death triggers, including enhanced defense responses, secondary metabolism and proteolysis. FB1 also led to accumulation of proteins involved in protein transport, whereas avrRxo1-triggered cell death led to accumulation of oxidative stress-responsive proteins. Similar to the transcriptome data, SA treatment resulted in the earliest changes on the proteome level, which largely remained the same over time and included mitochondrial processes and glutathione metabolism. Noteably, photosynthesis and other chloroplast processes were among the depleted pathways in all treatments from 48 hpi and onwards.

### Correlation of transcriptome and proteome reveals cell death markers

To better understand the mechanisms underlying cell death in maize leaves, we performed a correlative analysis of the transcriptomic and proteomic data. To increase our detection area, all DEGs without log_2_FC cut-off (12841) were used together with all previously identified DAPs (1702). Initially, the Pearson correlation was calculated among all mRNA and all protein samples. Our analysis indicated that the overall correlation was quite low (correlation<0.5, *P*-value<0.05), with the 24 hpi-mRNA avrRxo1 samples having the highest significant positive correlations (cor.=0.35, *P*- value<2.2e-16 with 48hpi-protein; cor.=0.38, *P*-value<2.2e-16 with 60hpi-protein and cor.=0.29, *P*-value<2.2e-16 with 72hpi-protein) (Fig. S2). To dig deeper into the actual genes / proteins hidden behind those low, yet significant, correlations, correlation modules were generated, with each module representing the first time point where differential gene expression or differential protein abundance was observed (for transcriptomics FDR<0.05 and proteomics *P*-value<0.05, without log_2_FC cut-off) (Fig. 6A). In that way, we identified 200 correlation modules, 56 of which corresponded to either no change in gene expression but protein abundance, or altered transcription without change on the protein level (Fig. 6A). Genes included in each correlation module are presented in Table S5. Special focus was given to the modules derived from significant positive correlations, based on the Pearson correlation coefficient (*P*-value<0.05, Fig. S2). Those modules included genes which were up-regulated and showed protein accumulation (Q1, quadrant I, Fig. 6A, Table S5), or were down-regulated and showed protein depletion (Q3, quadrant III, Fig. 6A, Table S5). More specifically, for avrRxo1 Q1: 12/24/48hpi-mRNA x 48/60/72hpi-protein and Q3: 12hpi-mRNA x 60hpi-protein, 24/48hpi-mRNA x 48/60/72hpi-protein; for FB1 Q1: 24hpi-mRNA x 24/48/60hpi-protein, 48hpi-mRNA x 60hpi-protein and Q3: 24hpi-mRNA x 24/48/60hpi-protein, 48hpi-mRNA x 60hpi-protein; for SA Q1: 12hpi-mRNA x 24/48/60hpi-protein, 24hpi-mRNA x 24/48hpi-protein and Q3: 12hpi-mRNA x 24/48/60hpi-protein, 24hpi-mRNA x 24/48hpi-protein (Fig. 6A). Those modules were then used for the identification of cell death markers exhibiting the same behavior on transcript and proteome level.

**Figure 6:**
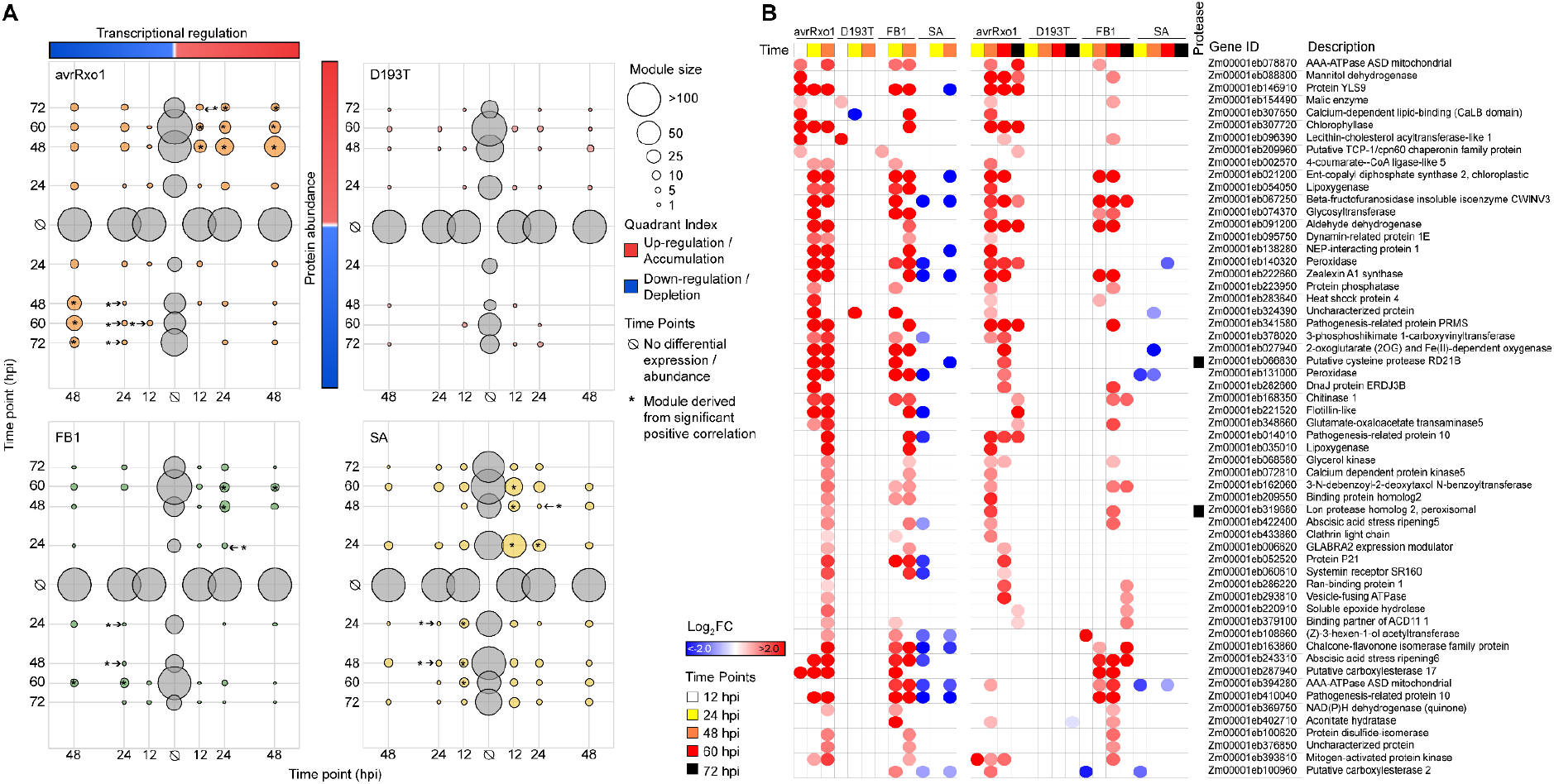
Identification of general cell death marker genes. (A) Correlation between gene expression and protein abundance. Correlation modules were created based on the first time point where differential gene expression and differential protein abundance was observed (for transcriptomics FDR<0.05 and proteomics P-value<0.05, without any log2FC cut-off). Each graph represents modules for one of the four treatments (avrRxo1, D193T, FB1 and SA) and each quadrant corresponds to a correlation between transcriptional regulation and protein abundance. As labelled with red and blue in the first graph (avrRxo1): upper left: gene down-regulation and protein accumulation, upper right: gene up-regulation and protein accumulation, down left: gene down-regulation and protein depletion, down right: gene up-regulation and protein depletion. Circles highlight the modules and their size reflects the number of genes/proteins in the modules. Asterisks mark modules derived from statistically significant positive correlation between mRNA and protein (Pearson correlation, P-value<0.05, for details see Figure S2). Genes from modules with statistically significant positive correlations were further analyzed for the identification of cell death markers. (B) Heatmap of selected avrRxo1- and FB1-driven common cell death marker genes. Markers were selected from the modules with statistically significant positive correlations between mRNA and protein, and they were picked for their specific gene expression/protein abundance in avrRxo1 and FB1. For a full list of markers see Table S5.

We found 58 general cell death markers, which were specifically present in avrRxo1 (three of them in avrRxo1 and D193T) and FB1 datasets and were either repressed or not differentially regulated in the SA datasets (Fig. 6B). Among those markers, two proteases were detected: a RD21B / CP1-like PLCP (Zm00001eb066830), and a serine-type peroxisomal Lon protease (Zm00001eb319680) (Fig. 6B, Table S6).

Among the identified 58 general cell death markers, we have tested four genes in independent experiments (Fig. S4). One of them, Zm00001eb287940, the putative carboxylesterase hsr203J was specifically induced only in response to the two cell death triggers, but not upon SA or JA treatment (Fig. S4 A-C). In the 1990s, hsr203J was already described as a HR marker in tobacco (Pontier et al., 1994, Pontier et al., 1998) suggesting that this gene might serve as a transcriptional cell death indicator in several plant lineages. The other three tested marker genes Zm00001eb307720 (chlorophyllase); Zm00001eb054050 (lipoxygenase 4) and Zm00001eb027940 (leucoanthocyanidin dioxygenase) were not only induced during cell death, but also in response to either both or only one of the immune-related hormones SA and JA. In response to SA the expression of Zm00001eb307720 was induced at 6 and 24 hpi and Zm00001eb054050 only at 6 hpi (Fig. S4 D-L). In the transcriptome analysis, although Zm00001eb307720 has shown up-regulation upon SA treatment at 12 and 24 hpi, this result was not statistically significant (Table S1). In addition, we did not anayze the transcriptome at 6 hpi, where the induction was more prominent according to qRT-PCR results (Fig. S4D).

Furthermore, we identified avrRxo1- and FB1-specific markers. Of these, 26 markers were specific for avrRxo1 function (not present in the D193T mutant), 9 markers were specific for the avrRxo1 protein (present in both, avrRxo1 and D193T) and 5 markers were explicit for FB1 (Fig. S3, Table S6). Among them, one putative aspartyl protease (Zm00001eb091230) specifically accumulating in response to the presence of avrRxo1 protein, was detected (Figure S5, Table S6).

### N-terminome analysis indicates changes in proteolytic activity during cell death

Various proteolytic activities have been implicated in the initiation and execution of cell death in plants (Salguero-Linares & Coll, 2019). To assess differential proteolytic activities during the course of RCD, we enriched N-terminal peptides by negative selection using HUNTER, followed by mass spectrometry-based identification (Weng et al., 2019). We set up two independent triplex dimethyl stable isotope labeling experiments, one comparing the NLR-induced cell death by avrRxo1 with the D193T and empty vector as control, and the other comparing FB1-treated samples with SA- and mock. In a first step, proteins were denatured and all primary amines differentially labelled using three different formaldehyde isotopologues for reductive demethylation. Samples were combined, digested with trypsin, and all non-N-terminal peptides modified at their trypsin-generated N-terminus with undecanal were removed. This step provides efficient simultaneous enrichment of N-terminal peptides derived from endogenously acetylated protein N-termini and *in vitro* dimethylated peptides, representing protein N-termini with free primary amines *in vivo*. Combined analysis of four biological replicates at four time points identified 1887 N-terminal peptides in FB1 - treated samples and 2090 N-terminal peptides in the avrRxo1-treated samples. Positional annotation revealed that in 35% and 38% of the identified N-terminal peptides in avrRxo1 and FB1 samples matched position 1 and 2 in their corresponding protein models, representing full-length proteins after synthesis with intact methionine and excised methionine, respectively (Fig. 7A). Another 30% (avrRxo1) and 33% (FB1) matched known or predicted signal peptides (SP), mitochondrial targeting sequences (mTP) or chloroplast transit peptides (cTP). Finally, 34% of the N-termini in avrRxo1 and 28% in FB1, matched to positions within protein models not annotated as canonical maturation sites, respectively, with dimethylated peptides likely derived from post-translational proteolytic processing.

**Figure 7:**
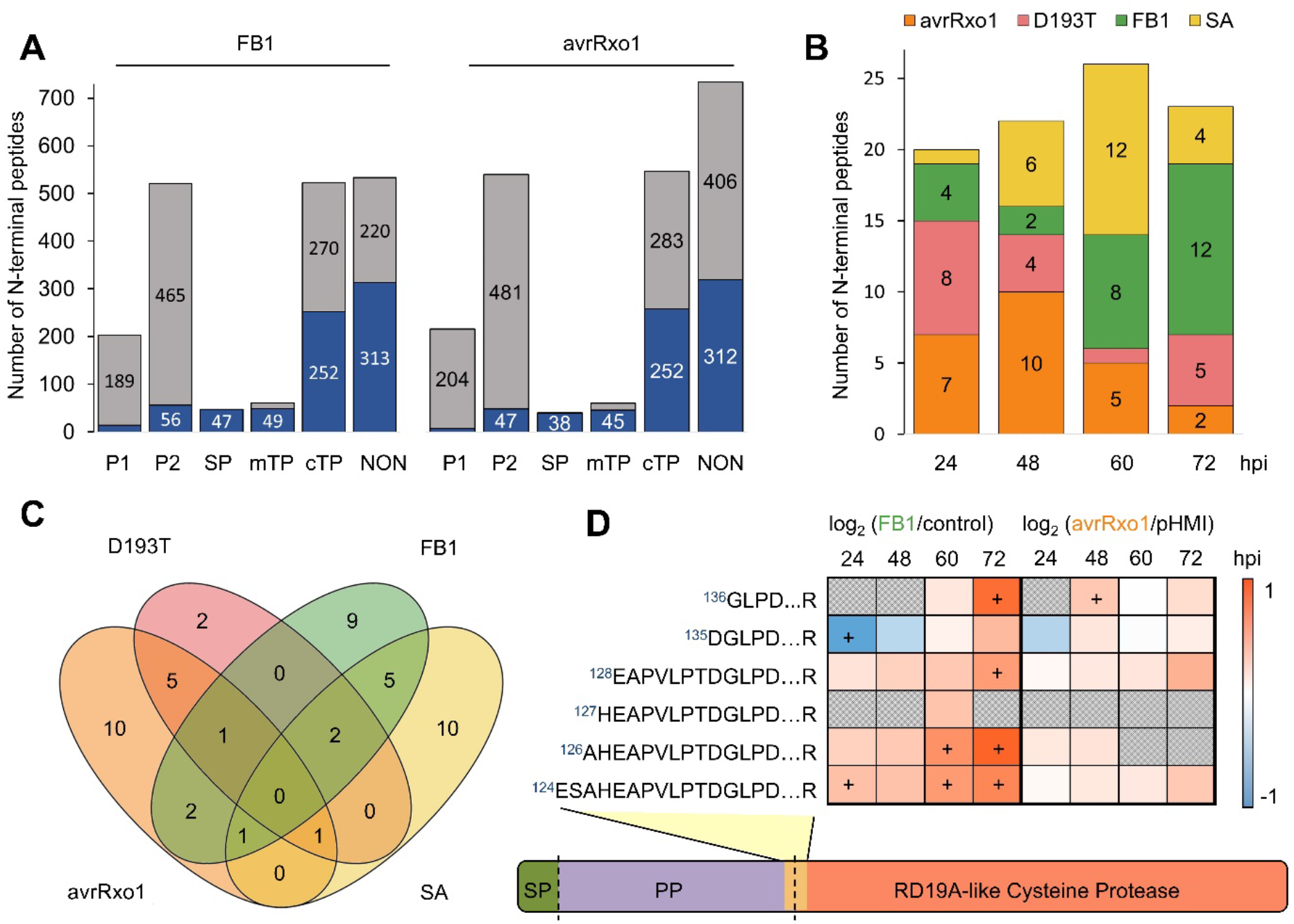
N-terminome analysis reveals likely activating cleavage of RD19A-like Papain-like Cysteine Protease in maize. (A) Positional annotation of the identified N-terminal peptides in each experiment. Grey, endogenously acetylated N-terminal peptide; blue: dimethylated, in vivo free N-terminal peptide; P1, position one in the protein model starting with Met; P2, position 2 in the protein model after methionine excision; SP, signal peptide; mTP/cTP, mitochondrial targeting sequence and chloroplast transit peptide; NON, non-canonical positions matching within the protein model. (B) Number of significantly accumulating (π-value > 1.1082, Xiao et al., 2014) non-canonical, dimethylated N-terminal peptides likely derived from proteolytic processing at each time point. (C) Venn diagram comparing the overlap of the proteolysis-derived N-terminal peptides significantly accumulating at any time point across conditions. (D) Schematic representation of N-terminal peptides mapping to the RD19A-like cysteine protease during cell death. Protein domain borders derived from UniProt annotation are indicated, peptide abundance is visualized as a heatmap with “+” indicating significant changes in abundance (π-value > 1.1082 or < −1.1082); Green box SP, signal peptide; Purple box PP, propeptide; grey-shaded fields in the heat map, no ratio determined.

To determine proteolytic cleavages altered during cell death, we next limited the dataset to N-terminal peptides quantified in at 3 replicates in at least one time point, resulting in 809 and 803 reliably quantified N-terminal peptides in the FB1 and avrRxo1-dataset, respectively (Tables S7 and S8). We then determined significant changes in N-termini abundance using a π-value > 1.1082 or <-1.1082 as cut off (Xiao et al., 2014). This approach identified 316 more and 362 less abundant N-terminal peptides at any time point or condition in the avrRxo1 and FB1 experiments, respectively (Fig. S5). Overall, the highest number of N-terminal peptides with significant changes were observed for avrRxo1 in the early time points (24 hpi and 48 hpi), with 67 significant changes in N-termini abundance at 24 hpi increasing to 80 at 48 hpi. This numbers drop at the later time points to 18 and 34 proteins. In the D193T, the FB1-treated and SA-treated samples a time-wise more stable significant protein abundance can be observed, with most changes in FB1 (ranging from 45 to 60 proteins) compared to SA (23 to 36 proteins) and D193T (20 to 43).

However, most of the N-terminal peptides with altered abundance mapped to positions expected from synthesis or known maturation and that reflected overall protein abundance. Only a very limited number of proteolysis-derived dimethylated, non-canonical N-termini accumulated, with peaks observed at 48 hpi after avrRxo1-triggered cell death and at 60 to 72 hpi for FB1-triggered cell death (Fig. 7B). Comparison of these accumulating proteolytic cleavages in each condition across all time points revealed two cleavages that were exclusively observed after both cell death triggers (Fig. 7C). One was a cleavage before Leu39 of an Acyl-Coenzyme A oxidase, just 5 amino acids downstream of an annotated peroxisomal transit sequence cleavage site in the homologous protein of *A. thaliana*. Therefore, the observed cleavage site is very likely also a peroxisomal targeting sequence cleavage in the maize protein (Fig. S5C) which reflects a change in mature protein abundance. Remarkably, the second common proteolysis-derived N-terminal peptide started at Gly136 of the RD19A-like cysteine protease (Zm00001eb240700), which is annotated as activating propeptide cleavage site in the UniProt database (indicated by dotted line in Fig. 7D). The dataset additionally contained five N-terminal peptides matching the same RD19A-like protease within the propeptide, which similarly accumulated significantly after FB1 treatment at 24, 60 and 72 hpi, suggesting intermediate activation steps and N-terminal ‘ragging’ by aminopeptidases (Fig. 7D). The same peptides corresponding to RD19A-like were also more abundant during avrRxo1-triggered cell death, however their accumulation was not significant (Fig. 7D). Similarly, we observed additional N-terminal peptides that may indicate altered activation of distinct proteases during defense responses: After SA treatment, an N-terminal peptide starting at Ala107 of CP1A (Zm00001eb068400), close to its predicted activation site, increased early at 24 hpi, whereas the same peptide appeared in significantly lower abundance in the presence of avrRxo1 and D193T. Besides, SA treatment also resulted in accumulation of peptides starting at Gly129 at the activation site of CP1B (Zm00001eb432170) (Fig. S5D). The activating cleavage observed for CP1A and CP1B is in agreement with their well-established activation after SA treatment in maize roots and shoots (van der Linde et al., 2012; Schulze Hyunk et al., 2019). In the AvrRxo1 data, we observed significant accumulation in N-terminal peptides starting at Gly119 of an aspartic protease (Zm00001eb116230), a homolog of the secreted aspartic proteases 1 and 2 (SAP1 and SAP2) in *A. thaliana*, and at Tyr34 of the mitochondrial serine protease CLPP (Fig. S5D). The cleavage site in the aspartic protease matches to a position that might indicate propeptide cleavage and activation (Soares et al., 2019). Interestingly, the *A. thaliana* homologues SAP1 and SAP2 were reported to contribute to antimicrobial resistance by cleavage of the secreted bacterial effector protease MucD (Wang et al., 2019). In contrast, the cleavage site in CLPP is likely a mitochondrial targeting sequence cleavage site and thus likely reflects abundance of the mature, imported protease (Fig. S5D). The CLPP N-terminal peptide appeared to accumulate also early after infiltration with FB1, but did not meet the significance criteria due to high variation among the replicates (Table S7). Thus, HUNTER N-terminome data provided a snapshot of altered protease activities and suggests that proteases, including a RD19A-like protease, are specifically activated in both NLR-mediated and toxin-induced cell death in maize.

## Discussion

We performed a multiomics analysis to determine components involved in NLR- and toxin-induced cell death in maize leaves. The combination of transcriptome, proteome and N-terminome analysis allowed us to identify shared and also unique markers for the Avr-R interaction and the toxin FB1-triggered cell death. A module correlation analysis of transcriptome and proteome data led to the identification of 58 general cell death markers, 5 FB1- and 26 avrRxo1-specific marker genes which may play a role as initiators or executers of cell death in maize. Based on GO Terms, no biological process could be associated to about a third (21 of the 58 genes / proteins) of the identified markers. Other markers were annotated as involved in pathways related to defense response, phosphorylation, abscisic-acid activated signaling, carbohydrate metabolism, gibberellin biosynthesis and lipid metabolism. The two phytohormones abscisic acid (ABA) and gibberellin (GA) participate in regulated cell death during plant development. Epidermal cell death of root cortex and node cells in rice is modulated, negatively and positively, by ABA and GA, respectively (Steffens & Sauter, 2005). Similarly, GA promotes cell death, while ABA inhibits cell death in the barley cereal aleurone layer (Fath et al., 2001). In maize, the appropriate onset and development of RCD during endosperm development is thought to be established via a balance of ABA and ethylene (Young & Gallie, 2000). Recently, susceptibility towards *Botrytis* infection in *A. thaliana* has been shown to be promoted by ABA suggesting a role of ABA in immune related cell death (Cui et al., 2019). The fact that we identified marker genes and proteins potentially involved in abscisic-acid activated signaling pathway and gibberellin biosynthetic process hint towards an involvement of both hormones in immune related cell death in maize. In accordance, our comparison of the avrRxo1- and FB1-triggered cell death with the phytohormone SA showed that the SA-induced response leads to a strong early transcriptional reprogramming and proteomic activation of immune-related components which decreases over time. This is in agreement with an early SA-responsive gene analysis showing that a large number of positive regulators of defense signaling are strongly up-regulated one hour after SA treatment (Ding et al., 2018). In our experiments SA alone did not trigger cell death but avrRxo1 and FB1 triggered first SA responses. We expected to uncouple phytohormonal responses from the initiation of cell death but this was not possible, most likely because these pathways are interconnected. SA has been shown to be required for spontaneous cell death in several lesion-mimic *A. thaliana* mutants (Radojičić et al., 2018) and to be involved in immunity against biotrophic pathogens, accumulating around infection sites in a concentration gradient where it triggers various defense responses (Enyedi et al., 1992; Dorey et al., 1997; Fu et al., 2012; Yan & Dong, 2014).

Enriched GO Terms identified for DEGs or DAPs in response to avrRxo1 or FB1 were mostly unique for the respective trigger, indicating that avrRxo1 and FB1 induce different cell death pathways in accordance with their different modes of action and suggesting a thigh regulation and control of the cell death process. Enriched GO terms being shared between avrRxo1 and FB1 include “defense and immune response”, “receptor signaling pathways”, “defense molecules” and “secondary metabolism”, indicating that a general plant immune response is commonly activated. Consequently, five general cell death markers are involved in phosphorylation processes including a MAPK, as well as a CDPK indicative for an activation of signaling cascades, similar to the SA pathway. This is not surprising in case of avrRxo1 triggered-cell, which was delivered by a bacterial pathogen that actually induces SA responses. For FB1, the situation is different, since the trigger was directly delivered through vacuum infiltration, but still it activated immune responses related to the SA signaling pathway. Recently, a spatio-temporal transcriptome analysis of HR in *A. thaliana* also identified various enriched processes including cellular response to hypoxia and defense response to bacterium (Salguero-Linares et al., 2022). Four of the 14 enriched processes found in the transcriptome data of cells undergoing cell death in *A. thaliana* were also enriched in maize upon avrRxo1 treatment, namely response to heat, defense response to fungus, response to SA and response to hydrogen peroxide (Salguero-Linares et al., 2022). This substantiates that NLR-mediated cell death responses are at least partially conserved between monocot and dicot species.

In recent literature, a couple of genes playing a role in plant RCD were identified. Two genes being crucial in the HR, triggered by the autoactive maize NLR gene Rp1-D21 (Murphree et al., 2020), were also found up-regulated in our analysis: i) HSP90 (Zm00001eb199590) expression was induced in response to avrRxo1 at 24 hpi ii) a hydroxycinnamoyl-CoA shikimate/quinate (Zm00001eb006410) which was induced upon FB1 and avrRxo1 treatments. Similar to Salguero-Linares and colleagues, we have identified AAA-ATPases, as well as one vesicle-fusing ATPase as cell death markers. The expression of an ATPase (AT3G63380) was also reported to be upregulated in a meta-transcriptome analysis upon biotic stresses (Olvera-Carrillo et al., 2015). AAA-ATPases play a role in various processes including proteolysis, DNA replication and membrane fusion (Snider et al., 2008). In the last years, AAA-ATPases were reported to be involved in immunity and cell death. *A. thaliana* overexpression lines of the mitochondrial outer membrane AAA-ATPase AtOM66 show enhanced cell death and are more resistant towards *Pseudomonas syringae* (Zhang et al., 2014). Contrary, suppression of *Nt*AAA1 in tobacco resulted in an increased HR response and less bacterial growth of *P. syringae* pv. *glycine* (Lee & Sano, 2007). The multivesicular body (MVB) associated AAA-ATPase LRD6-6 from rice was reported to be required for suppression of cell death and plant immunity (Zhu et al., 2016). In conclusion, AAA-ATPases can exhibit inhibitory or accelerating effects on cell death and their function in immune related cell death of maize remains to be investigated. Another known cell death marker, the serine hydrolase hsr203J (Zm00001eb287940) was also identified and independently verified in our analysis on both, transcript and protein levels. In 1994, hsr203J was first discovered as cell death marker in tobacco plants where it was shown to be induced upon infection with *Ralstonia solanacearum* (Pontier et al., 1994). In tomato, the expression of a homologous hsr203j gene is induced upon recognition of AVR9 of *Cladosporium fulvum* (Pontier et al., 1998). The expression of a putative hsr203J-like protein in grapevine was enhanced in response to the necrotrophic pathogen *Botrytis cinerea* (Bézier et al., 2002). In a transcriptome meta-analysis, an alpha / beta hydrolase superfamily protein (AT1G68620), which is one of the closest orthologues of tobacco hsr203j, was also found to be commonly upregulated in response to biotic stresses in *A. thaliana* (Olvera-Carrillo et al., 2015). The fact that hsr203j is induced in several plant lineages during HR, suggests that this serine hydrolase is an important, broadly conserved component of the immune-related cell death in plants.

In mammals, cysteine proteases of the caspase family initiate and execute apoptosis and regulate pro-inflammatory RCD (Bateman et al., 2021). Likewise, proteolytic activities are expected to play a significant role in plant RCD, potentially as triggers, but certainly during execution and corpse clearance (Huysmans et al., 2017). Early evidence suggested a role of caspase-like activities in plant RCD (reviewed in Bonneau et al., 2008), but plants do not have caspases (Minina et al., 2020) and to date no single protease family appears to be universally required for plant RCD (Salguero-Linares & Coll, 2019). Instead, a whole variety of different proteases belonging to the cysteine, serine and threonine families, some displaying caspase-like activities whereas others do not, have been described to be involved in HR and other forms of RCD (Sueldo & van der Hoorn, 2017; Balakireva & Zamyatnin, 2019; Salguero-Linares & Coll, 2019).

Interestingly, several proteases were found as cell death markers in our analysis. A peroxisomal ATP-dependent serine Lon protease (Zm00001eb319680) and CP1-like cysteine protease related to the *A. thaliana* RD21 (Zm00001eb066830) were identified as general markers with higher expression and higher accumulation after FB1- and arvRxo1-triggered cell death. RD21 of *A. thaliana* has previously reported to be involved in immunity, however the exact role has not been yet elucidated. On the one hand *A. thaliana* mutants lacking RD21 show an enhanced susceptibility towards *Botrytis cinerea* (Shindo et al., 2012), while on the other hand lack of RD21 led to enhanced resistance to *B. cinerea* in detached leaf assays (Lampl et al., 2013). However, increased transcript and protein abundance is not a requirement for increased activity during RCD, as for example PLCPs are regulated by proteolytic prodomain cleavage (Misas-Villamil et al., 2016). Our N-terminome dataset suggests such post-translation regulation of PLCP activity during maize RCD, since we observed accumulation of a specific N-terminal peptide at Gly136 of a RD19A-like cysteine protease (Zm00001eb240700), indicating activation of this protease during both, FB1 toxin-induced and avrRxo1-triggered cell death. This RD19A-like cysteine protease was previously also shown to be activated in the apoplastic space of roots treated with SA (Schulze Hüynck et al., 2019). In avrRxo1-triggered samples, a modest but significant increase of this N-terminal peptide is seen after 48 hpi, the time when first clear cell death symptoms are observed. Consistently, the same peptide accumulates strongly at 60 and 72 hpi after FB1 treatment, when cell death appears. Remarkably, this protease was not found as upregulated or more abundant during the course of our study suggesting that post-translational activation of the available pro-protease pool might contribute to the cell death phenotype. Despite evidence for regulated protease activities, we did not observe accumulation of proteolytic substrate cleavage products in our N-terminome analysis. This likely results from technical limitations and suggests to generate cell-compartment specific datasets in future experiments, which could increase depth and resolution of the N-terminome analysis. Also, activating proteolytic cleavages likely occur only at a specific time during the execution of the RCD program and cleaved substrates may accumulate only transiently. Nevertheless, our N-terminome data reinforced an active role for cysteine proteases, in particular PLCPs, in immune-related RCD in plants, although their mechanism and function need to be further elucidated.

In this study, we provide time-resolved, multi-omics information on biological processes regulated by effector- and toxin induced RCD as well as SA signalling in maize leaves. We identified several general and trigger-specific cell death markers that were consistently upregulated on the transcript level and accumulate also on the protein level. N-terminome analysis identified RD19A as a cysteine protease which might be directly involved in both immune-triggered cell death processes. Taken together, our data will provide a valuable resource for identification of common elements with conserved functions across distinct pathosystems and in other, developmentally controlled cell death programs across plant species. Future work will establish the function of the identified markers and dissect which of the activated processes are directly related to the execution of cell death, and which are indirect consequences of signals send from dying cells to the surrounding tissue.

## Material and Methods

### Transformation of *Xanthomonas oryzae* pv *oryzae* PXO99A

*Xanthomonas oryzae* pv *oryzae* PXO99A bacteria were grown on peptone-sucrose agar for 2-3 days at 28°C. Bacterial cells were resuspended in sterile water, collected in a reaction tube and washed three times with cold and sterile water. Approximately 500 ng plasmid DNA were added to 50 μl bacterial cell suspension. *Xanthomonas oryzae* pv *oryzae* PXO99A was transformed with plasmids listed in table S9. The cells were transformed using electroporation (1500 V for about 5 ms). After electroporation, 750 μl peptone sucrose medium was added to the cells and incubated for 3 h at 28°C at 220 rpm. Cells were plated on peptone-surose agar with the respective antibiotic and incubated for 3-7 days at 28°C.

### Treatment of maize leaves

For all experiments, maize plants (*Zea mays* B73) were grown on soil in a walk in phytochamber with a light period of 12 h with 1 h twilight at 28°C and a dark period of 7 h with 1 h twilight at 22°C. The second leaf of 8-9 day old plants was used for each experiment. Leaves were excised and vacuum infiltrated (5 repetitions of 240 mbar for 10 min, followed by ATM for 2 min) with a control solution (1% Dimethyl sulfoxide (DMSO) in H_2_O), salicylic acid (2 mM in 1% DMSO in H_2_O), Fumonisin B1 (50 μM in 1% DMSO), methyljasmonate (1 mM in 1% DMSO) or *Xanthomonas oryzae* pv *oryzae* PXO99A carrying pHMI, pHMI-avrRxo1 or pHMI-avrRxo1 D193T (OD = 0.002 in sterilized H_2_O). Samples were taken at 12, 24, 48, 60 and 72 hpi. Per sample, treatment and time point four whole leaves were pooled for each of four biological replicates.

### Detection and Quantification of Cell Death

The green light signal of the infiltrated leaves was detected using the blue epi-illumination source and a 530/28 nm filter from the Chemi-Doc MP System (Bio-Rad, Hercules, CA, USA). The quantification of the intensity of the green light signal was performed using the ImageLab software (Bio-Rad, Hercules, California, USA). Therefore, the leaf area of each individual leaf was selected using the freehand volume tool. Per time point, the background signal from one part of one leaf infiltrated with the respective control solution showing no cell death was subtracted from all leaves. The signal intensity was normalized to the leaf area (mm^2^).

### Extraction of RNA and RNA sequencing

Pooled maize leaves were homogenized in liquid nitrogen using mortar and pestle. For total RNA isolation, TRIzol™ reagent (Invitrogen, Darmstadt,Germany) was used according to the manufacturer’s instructions. 1 ml TRIzol™ was added immediately to approximately 100 mg plant material. To exclude DNA contamination, the Turbo DNA-Free™ Kit from Ambion (Ambion Life Technologies™, Carlsbad, USA) was used according to the manufacturer’s instructions. Sequencing library preparation and Illumina sequencing of mRNA was performed with 150-bp paired end reads at Novogene Bioinformatics Technology Co., Ltd. on an Illumina NovaSeq 6000.

### RNAseq data analysis

Paired-end RNAseq reads were quality trimmed using Trimmomatic, selecting for per-base quality score in the start and end of the reads ≥ 20. Subsequently, reads were mapped to the *Z. mays* genome (B73 reference genome version 5.0) with bowtie2 in very-sensitive end-to-end mode (Langmead & Salzberg, 2012). Successfully mapped reads were sorted with samtools and counted with HTseq-count (Anders et al., 2015) using intersection-nonempty mode for unstranded reads. Statistical analysis of differentially regulated genes between the treatments and the respective control samples was performed with edgeR (Robinson et al., 2009), using a quasi-likelihood negative binomial generalized log-linear model to fit the count data. Genes with log_2_FC>1 and FDR<0.05 were considered as significant differentially expressed genes. The MDS plot was generated using the Glimma R package. Heatmaps were constructed using log_2_ transformed fold-changes. Hierarchical clustering of the heatmaps was performed using 1 minus the Pearson correlation coefficient, leading to the identification of distinct expression modules. A higher stringency of log_2_FC>2 was only used for the hierarchical clustering, in order to increase the sensitivity of the clustering method. For every identified module, the average expression level for each treatment was calculated, and the Pearson correlation to the cell death readout of maize leaves was calculated. Correlations were considered significant if absolute correlation coefficient>0.5 and *P*-value<0.05. Raw data are publicly accessible in the NCBI Gene Expression Omnibus (Edgar et al., 2002) with accession number GSE220941.

### Gene Expression Analysis

2 μg DNase treated RNA per 20 μl reaction were used to generate cDNA with the RevertAid H Minus First Strand cDNA Synthesis Kit (Thermo Fisher Scientific, Waltham, USA) according to the manufacturer’s instructions. For gene expression analysis, 6.5 μl of 1:100 diluted cDNA per reaction were used with the GoTaq® qPCR Mastermix (Promega GmbH, Madison, USA) in a total volume of 15 μl. All qRT PCRs were performed in an iCycler system (Bio-Rad, Hercules, USA) with the following conditions: 95°C for 2 min, 45 cycles of 95°C for 30 s, 61°C for 30 s and 72°C for 30 s followed by fluorescence reading. For normalization, *GAPDH* was amplified in parallel on each plate. To exclude contaminations and primer-dimer formations, melting curves and no template controls were included and analyzed on each plate. Used Primers are listed in Table S10.

### Protein extraction and proteome sample preparation

The plant material was ground into a fine powder using mortar and pestle under liquid nitrogen. Approximately 1 mL of powder was transferred to a 2 mL reaction tube, dissolved in 1.5 mL of a denaturing buffer (6 M Gua-HCl, 100 mM HEPES, adjusted to pH 7.4 and supplemented with 1x HALT protease inhibitor cocktail (Thermo Fisher Scientific, Rockford, USA) and vortexed. Afterwards the sample was filtered with Miracloth (Merck KGaA, Darmstadt, Germany) and the flowthrough collected in a 2 mL reaction tube. The protein concentration was measured by using the Pierce BCA Protein Assay Kit (Thermo Fisher Scientific, Rockford, USA). For proteome analysis, 20 μg proteome and treated with 0.5μL Benzonase (~26 U/μL). 10mM DTT was added to the samples and incubated shaking at 37°C for 30 min, cooled to RT, 50mM Chloroacetamide (CAA) were added and incubated in the dark for 30min. The reaction was quenched by adding additional 50 mM DTT and incubating for another 20min before protein purification using SP3-beads (Cytiva Europe GmbH, Freiburg im Breisgau, Germany) as described in Hughes et al., 2019. The samples were resuspended in trypsin digestion buffer (volume accordingly to a final protein concentration of 2μg/μL) containing 100 mM HEPES and 2.5 mM CaCl2 adjusted to pH 7.4 and digested with trypsin (1:100 trypsin: proteome) over night at 37°C. The next day the samples were desalted by using self-made STAGE tips packed with C18-material (Rappsilber et al., 2007). The samples were resuspended in 0.1% formic acid.

### HUNTER N-termini enrichment

For the HUNTER N-terminome analysis, 100 μg extracted protein sample were treated with 1 μL Benzonase at 37°C (~26 U/μL) for 30 min, 10 mM DTT added and incubated shaking at 37°C for 30 min, followed by cooling to RT, addition of 50 mM CAA and incubation in the dark for 30 min. The reaction was quenched by adding an additional 50 mM DTT and incubated for another 20 min before protein purification using SP3-beads as described (Mikulášek et al., 2021). Protein samples were resupended in denaturing buffer (6 M Gua-HCl, 100 mM HEPES, pH 7.4) and stable isotope labelled using with 30 mM ^12^CH_2_O and 30 mM sodium cyanoborohydride (light), 30 mM 12CD_2_O and 30 mM sodium cyanoborohydride (medium) or 30 mM 13CD_2_O and 30 mM sodium cyanoborodeuteride (see Table S11 for labelling scheme) and incubated for 2 h at 37°C. The reactions were quenched by adding 1 M Tris-HCl pH 6.8 and incubating for 30 min at 37°C. The samples were combined into two triplex experiments (control+SA+FB1 and pHMI+D193T+avrRxo1, respectively), followed by protein purification with SP3-beads before, resuspension in digestion buffer and overnight digestion with trypsin as descibed for proteome samples. The next day, primary amines of trypsin-generated peptides were tagged with the hydrophobic aldehyde undecanal (UD). For this 3.3 μg UD in 40% EtOH per 100 μg peptides and freshly prepared sodium cyanoborohydride (final 30 mM) were added and incubated for 1 h at 37°C. Afterwards, undecanal-tagged peptides and excess reagents were removed by passing the reaction through HRX-reverse phase spin-columns (Macherey-Nagel GmbH & Co, KG, Dueren, Germany). The flow-through containing the enriched N-termini were dried with a SpeedVac to a remaining volume of ~50 μL. Afterwards the samples were desalted by using self-made STAGE tips packed with C18-material (Rappsilber et al., 2007). The samples were resuspended in 0.1% formic acid.

### Mass spectrometry data acquisition

The LC-MS/MS analysis was performed with an UltiMate 3000 RSCL nano-HPLC system (Thermo Fisher Scientific Inc., Waltham, USA) coupled to an Impact II Q-TOF mass spectrometer (Bruker) using a CaptiveSpray ion source (Bruker Daltonik GmbH, Bremen, Germany) operated with an ACN-saturated nitrogen gas stream was used. The peptides were loaded on an μPAC pillar array trap column (1 cm length, PharmaFluidics, PharmaFluidics, Ghent, Belgium) and separated on a μPAC pillar array analytical column (50 cm flowpath, PharmaFluidics, Ghent, Belgium). Peptides were eluted using a 2 h elution protocol that included an 80 min separation gradient from 5% to 35% solvent B (solvent A: H_2_O+0.1% FA, solvent B: ACN+0.1% FA) at a flow rate of 600 nl min^−1^, with columns temperated at 40°C. Proteome data were acquired using a Data Independent Acquisition (DIA) method with 400 to 1200 precursor mass windows for MS1, with 32 windows of 25 m/z size with 0.5 m/z overlap and 200 to 1750 for MS2 at 15 Hz. HUNTER N-terminome experiments were analysed in data-dependent acquisition (DDA) mode: MS spectra were acquired at 5 Hz in m/z range from 200 to 1400, with the 14 most intense precursors selected for fragmentation at 5 to 20 Hz in an intensity-dependent manner. Due to technical factors, the first steps of the analysis were conducted independently for the two experiments avrRxo1/D193T/pHMI and FB1/SA/control.

### Proteome data analysis

For downstream analysis, datasets from raw intensities were combined for the two experiments (avrRxo1/D193T/pHMI and FB1/SA/control), then only unique proteins detected in at least three out of four replicates in at least one condition were considered valid. Initially, all missing values (NA) were replaced with zeros. Thereafter, raw intensities were subjected to per-row (per-peptide) scaling normalization between 0 and 1. Subsequently, all 0 values were replaced with 1e-10 for mathematical reasons, to avoid error upon fold-change calculations, and the log_2_FC of the treatments relative to the respective controls was calculated. Additionally, the log_2_FC based on the raw intensities was calculated (labelled as log_2_FC_R). For statistics, Student’s t-test was performed per protein, for each tested comparison and only those with *P*-value<0.05 were considered as significant differentially abundant proteins. The MDS plots were generated using the Glimma R package with raw data. Heatmaps were constructed using log_2_ transformed fold-changes derived from normalized data, and the hierarchical clustering was performed using 1 minus the Pearson correlation coefficient, leading to the identification of distinct protein abundance modules. For each module, the average protein abundance for each treatment was calculated, and the Pearson correlation to the cell death readout of maize leaves was calculated. Correlations were considered significant if absolute correlation coefficient>0.5 and *P*-value<0.05.

### N-terminome data analysis

For peptide to sequence matching of the mass spectrometry data, we generated a custom protein database file based by filtering the maizeGDB database (B73 reference genome version 5.0) with only proteins observed in the RNA-seq dataset, combined with a contaminant list. DIA identification of peptides was performed with the software DIA-NN (version 1.8) in double-pass mode with a precursor FDR of 1%.

For HUNTER data, peptides were identified by matching spectra against the database using the MaxQuant software version 2.0.3.0 (Tyanova et al., 2016). We used standard settings, if not mentioned otherwise. ‘Type’ was set to three labels: Light (+28.0313), medium (+32.0564) and heavy (+37.0756) dimethylation of Lys residues. Peptide N-terminal acetylation, Met oxidation and pyro-Glu were set as variable modifications, and enzyme specificity to semi-specific ‘ArgC’. Furthermore, the ‘requantify’ option was enabled and the minimum score for recalibration was set to 47. N-terminome data were further processed and annotated with the MANTI script (version 5.0) (Demir et al., 2021) in combination with subcellular localization predictions by targetP 2.0 (Armenteros et al., 2019).

### Module generation of co-expressed genes or co-abundant proteins

Heatmaps of transcriptomic and proteomic data were constructed using log_2_ transformed fold-changes derived from normalized data, and the hierarchical clustering was performed using 1 minus the Pearson correlation coefficient, leading to the identification of distinct gene expression or protein abundance modules. For each module, the average gene expression/protein abundance for each treatment was calculated, and the Pearson correlation to the cell death readout of maize leaves was calculated. Correlations were considered significant if absolute correlation>0.5 and *P*-value<0.05.

### GO enrichment analysis

GO enrichment analysis was performed with ShinyGO (version 0.76.3; Ge et al., 2020), with *P*-value cut-off 0.05, using Fisher’s exact test. Networks of GO term relationships were constructed in R, followed by manual curation of the network clusters to highlight the representative enriched pathways.

### Correlation between transcriptome and proteome

Correlation modules between transcriptome and proteome were created based on the first time point where differential gene expression and differential protein abundance was observed. More specifically, for transcriptomics FDR < 0.05 and proteomics *P*- value < 0.05, without any log_2_FC cut-offs, were used as inputs. Eventually, each module represented one of the following: gene up-regulation/protein accumulation, gene up-regulation/protein depletion, gene down-regulation/protein accumulation, gene down-regulation/protein depletion, for all the tested time points. Pearson correlation between transcriptomic and proteomic datasets was calculated and modules which derived from significant correlations (*P*-value < 0.05) were used for the identification of cell death markers.

## Acknowledgements

We thank Prof. Wolf B. Frommer for providing the bacterial strain *Xanthomonas oryzae* pv. *oryzicola* PXO99A and Prof. Jan E. Leach for providing the plasmids used in this study. This work was supported by Deutsche Forschungsgemeinschaft (DFG, German Research Foundation, Project-ID 414786233 – SFB 1403) and from the Cluster of Excellence on Plant Sciences (CEPLAS) funded by the Deutsche Forschungsgemeinschaft (DFG, German Research Foundation) under Germany’s Excellence Strategy – EXC 2048/1 – project ID: 390686111.

## Author Contribution Statement

GD, PFH and JCMV supervised and designed the project. GD, PFH, JCMV and SB conceived and designed the experiments. SB, JCM and UM conducted all experiments except for MS and HUNTER that were performed by MM. GS did the bioinformatic analysis of the RNASeq and MS data, MM analyzed the HUNTER data. All authors discussed and interpreted the data. SB and GS wrote the manuscript with contributions from all authors.

## Supplemental information

**Figure S1:** Detailed cell death development in maize.

**Figure S2:** Correlation between transcriptome and proteome in cell death of maize leaves.

**Figure S3:** Selection of avrRxo1- and FB1-specific cell death marker genes.

**Figure S4:** Verification of cell death marker genes for FB1- and avrRxo1-induced cell death.

**Figure S5:** Protein N-terminal peptides with altered abundance during maize RCD and positional mapping of selected protein N-termini.

**Table S1:** List of transcriptomics data (gene expression).

**Table S2:** List of enriched GO terms in the transcriptomic dataset.

**Table S3:** List of proteomic data (normalized and non-normalized FCs).

**Table S4:** List of enriched GO terms in the proteomic dataset.

**Table S5:** All markers (specific and general) in detail.

**Table S6:** List of N-terminal peptides

**Table S7:** N-terminal peptides identified by HUNTER N-terminome analysis after infiltration of fumonisin B1

**Table S8:** N-terminal peptides identified by HUNTER N-terminome analysis after infiltration of Xanthomonas oryzae expressing the effector avrRxo1

**Table S9:** Table of Plasmids used to transform *Xanthomonas oryzae* pv *oryzae* PXO99A

**Table S10:** Oligonucleotides used in this study.

**Table S11**: Labelling scheme for HUNTER

