## Supplementary Figures for "Combination of transcriptomic, proteomic and degradomic profiling reveals common and distinct patterns of pathogen-induced cell death in maize"

### Supplemental Figures

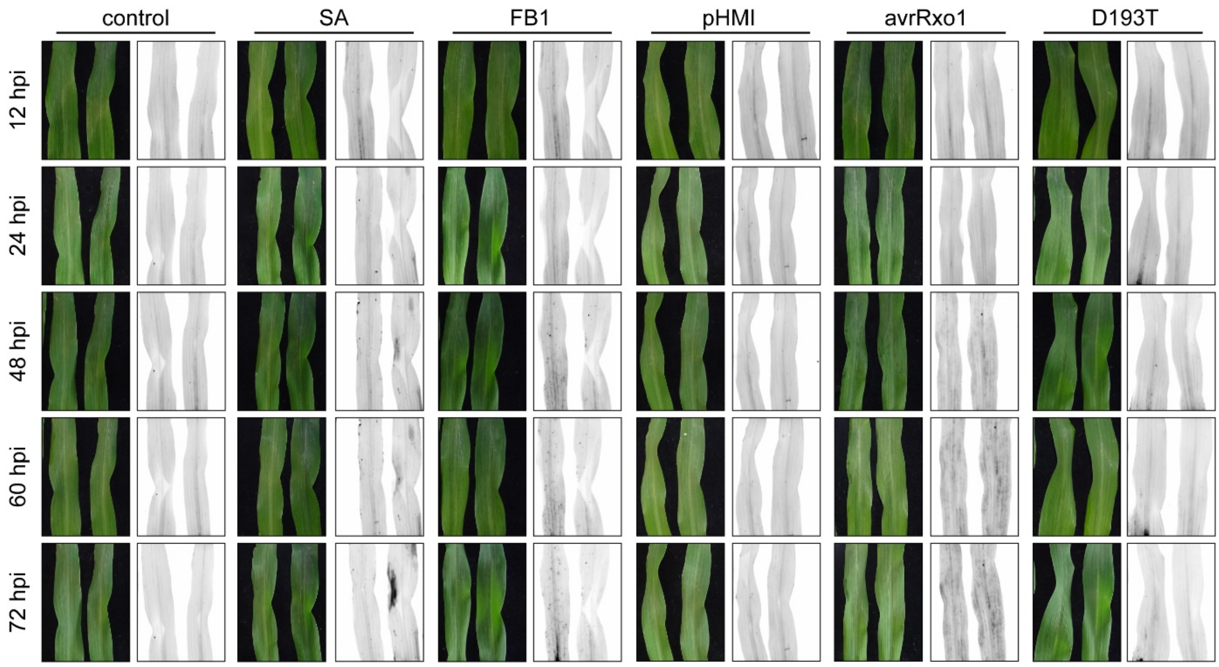


**Figure S1: Detailed cell death development in maize.** Maize leaves were vacuum-infiltrated with a control solution, SA [2 mM], FB1 [50 µM] or *Xanthomonas oryzae* pv. *oryzae* PXO99A carrying either the empty vector (pHMI), avrRxo1 or avrRxo1 D193T (D193T) at an OD=0.02. Cell death development was observed by taking colorimetric pictures and detection of green light emission at 12, 24, 48, 60 and 72 hpi.

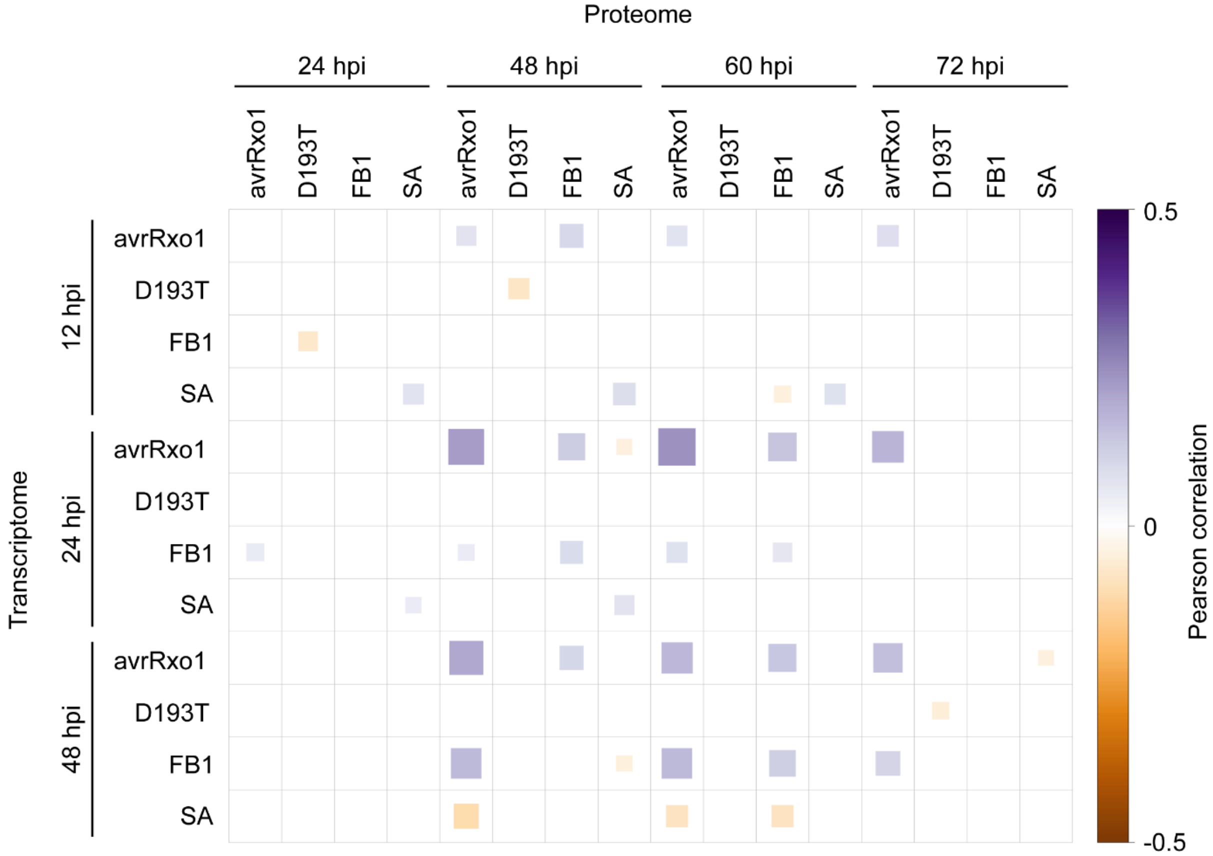


**Figure S2: Correlation between transcriptome and proteome in cell death of maize leaves.** Pearson correlation reflecting the weak correlation observed between transcriptomic and proteomic responses. Statistically significant correlations (*P*-value<0.05) were used in Fig. 6 to pick correlation modules for cell death marker selection.

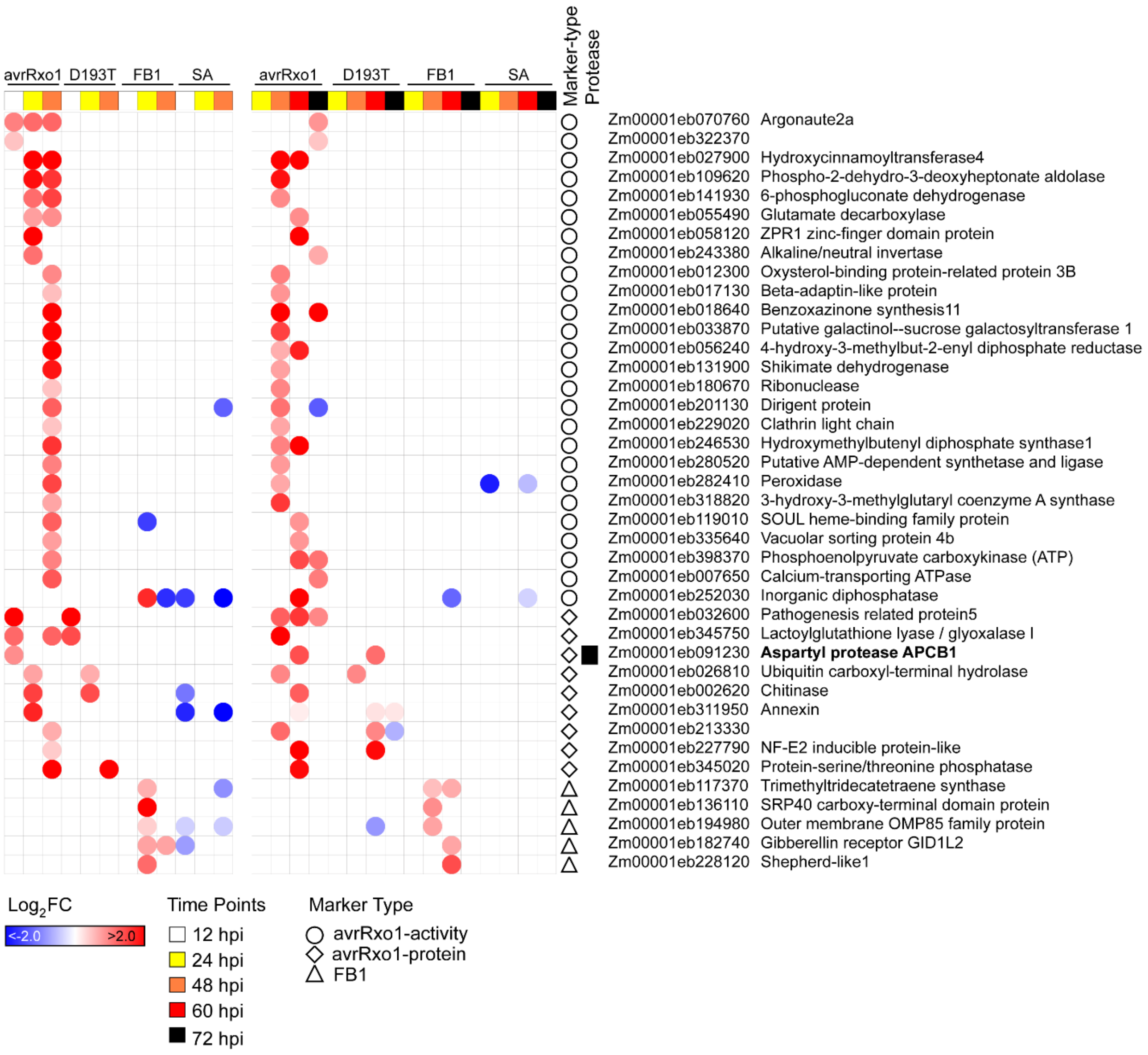


**Figure S3: Selection of avrRxo1- and FB1-specific cell death marker genes.** Markers were selected from the modules which showed significant correlations between mRNA and protein (see Fig. 6A), and they were picked for their specific gene expression/protein abundance in avrRxo1 (labelled as avrRxo1-kinase/cell death activity markers), avrRxo1 and D193T (labelled as avrRxo1-protein markers) or FB1 (labelled as FB1 markers).

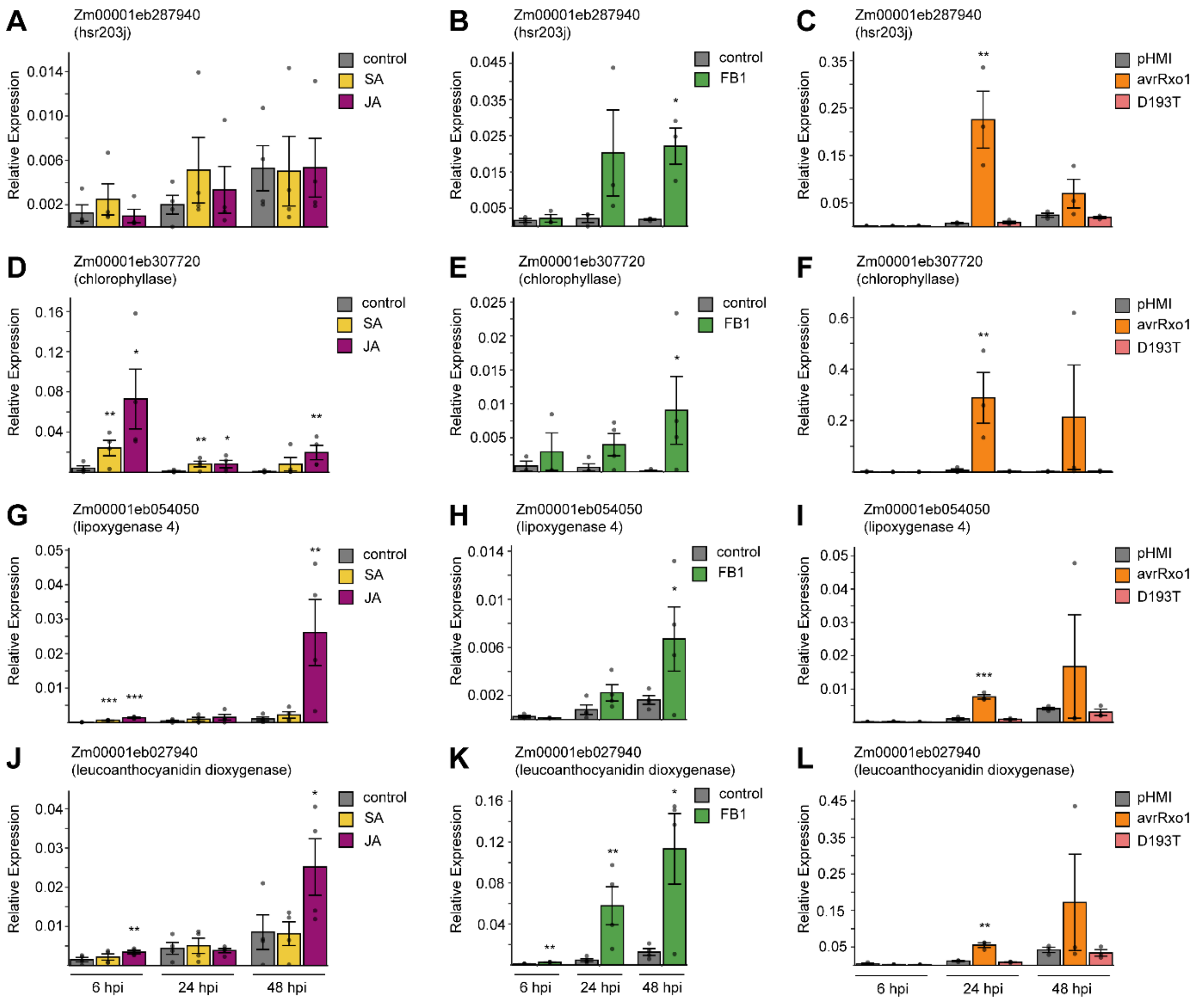


**Figure S4: Verification of cell death marker genes for FB1- and avrRxo1-induced cell death.** qRT-PCR data of four identified cell death marker (A-C) *Zm00001eb287940* (hsr203j), (D-F) Zm00001eb307720 (chlorophyllase), (G-I) Zm00001eb054050 (lipoxygenase 4) and (J-L) Zm00001eb027940 (leucoanthocyanidin dioxygenase). Maize leaves were treated with control solution (1% DMSO), SA [2 mM in 1% DMSO], JA [1 mM in 1% DMSO], FB1 [50 µM in 1% DMSO] or *Xanthomonas oryzae* pv. oryzae PXO99A carrying either the empty vector (pHMI), avrRxo1 or avrRxo1 D193T (D193T) with $OD_{600}$=0.02 and samples were taken at the indicated time points. Graphs show relative expression to the housekeeping gene *GAPDH*. Error bars represent standard error of mean of three to four independent biological replicates. Statistical significance was verified with unpaired student's t-test with: ns = p > 0.1, * = p < 0.1, ** = p < 0.05 and *** = p < 0.001.

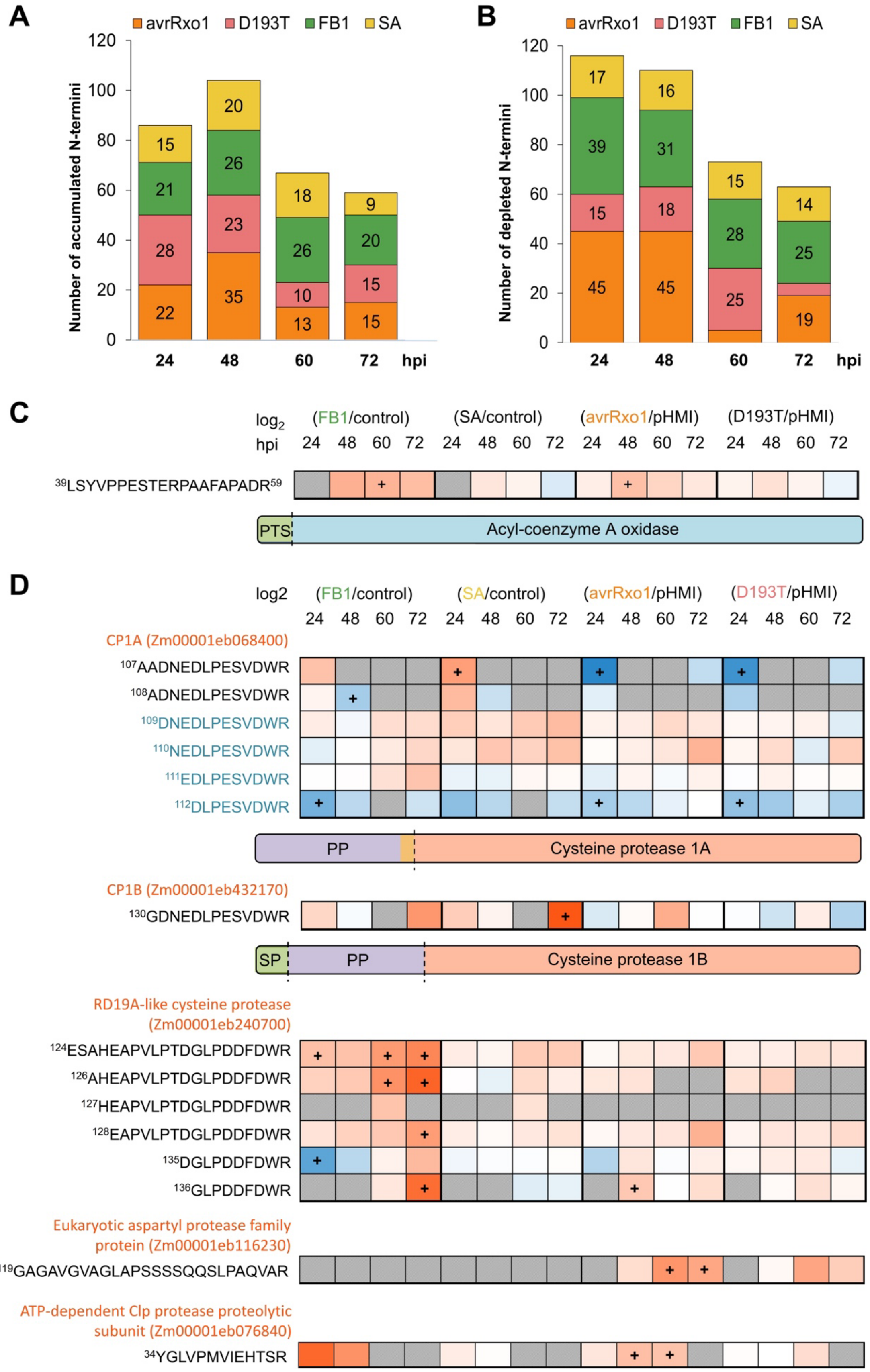
A
avrRxo1 D193T FB1 SA
Number of accumulated N-termini
0 20 40 60 80 100 120
24 48 60 72 hpi
22 28 21 15
35 23 26 20
13 10 26 18
15 15 20 9
B
avrRxo1 D193T FB1 SA
Number of depleted N-termini
0 20 40 60 80 100 120
24 48 60 72 hpi
45 15 39 17
45 18 31 16
25 28 15
19 25 14
C
log2 (FB1/control) (SA/control) (avrRxo1/pHMI) (D193T/pHMI)
hpi 24 48 60 72 24 48 60 72 24 48 60 72 24 48 60 72
39LSYVPPESTERPAAFAPADR59
PTS
Acyl-coenzyme A oxidase
D
log2 (FB1/control) (SA/control) (avrRxo1/pHMI) (D193T/pHMI)
24 48 60 72 24 48 60 72 24 48 60 72 24 48 60 72
CP1A (Zm00001eb068400)
107AADNEDLPESVDWR
108ADNEDLPESVDWR
109DNEDLPESVDWR
110NEDLPESVDWR
111EDLPESVDWR
112DLPESVDWR
PP
Cysteine protease 1A
CP1B (Zm00001eb432170)
130GDNEDLPESVDWR
SP
PP
Cysteine protease 1B
RD19A-like cysteine protease (Zm00001eb240700)
124ESAHEAPVLPTDGLPDDFDWR
126AHEAPVLPTDGLPDDFDWR
127HEAPVLPTDGLPDDFDWR
128EAPVLPTDGLPDDFDWR
135DGLPDDFDWR
136GLPDDFDWR
Eukaryotic aspartyl protease family protein (Zm00001eb116230)
119GAGAVGVAGLAPSSSSQQSLPAQVAR
ATP-dependent Clp protease proteolytic subunit (Zm00001eb076840)
34YGLVPMVIEHTSR

**Figure S5: Protein N-terminal peptides with altered abundance during maize RCD and positional mapping of selected protein N-termini.** Number of N-terminal peptides (A) significantly accumulating (π-value > 1.1082) or (B) significantly depleted (π-value < -1.1082) at each time point, irrespective of their positional annotation. Positional mapping of selected proteolysis-generated, dimethylated N-terminal peptides (C) derived from Acetyl-CoA- oxidase significantly accumulating specifically after both cell death triggers, (D) from maize proteases, the papain-like cysteine proteases CP1A and CP1B (note that peptides indicated in blue cannot be distinguished from CP1B) , for RD19A-like cysteine protease, for the aspartyl protease and for mitochondrial CLPP. Protein domain borders derived from UniProt annotation are indicated, green box SP, signal peptide; Purple box PP, propeptide; peptides abundances are visualized as heatmaps with, "+" indicating significant changes in abundance (π-value > 1.1082 or < -1.1082) and grey-shaded fields indicate conditions were no peptide was not observed.

.
